# Redeployment of Archaeal ESCRT Systems Reveals Conserved Mechanisms Underlying Eukaryotic Complexity

**DOI:** 10.1101/2025.11.01.685982

**Authors:** Kishore Babu Naripogu, Yee Han Tee, Yosuke Senju, Haokun Li, Alexander Bershadsky, Tomoyuki Honda, Robert C. Robinson

## Abstract

Eukaryotic ESCRT complexes remodel membranes at the plasma membrane and internal compartments, yet how these systems evolved from prokaryotic precursors functioning at the cellular membrane remains unresolved. Here, we show that ESCRT-II, ESCRT-III, VPS4, and ubiquitin from *Candidatus Prometheoarchaeum syntrophicum* (MK-D1), an Asgard archaeon closely related to eukaryotes, are recruited to canonical ESCRT sites in human cells, including centrosomes, midbodies, nuclear envelopes, endosomes despite the absence of these structures from MK-D1 morphology. In addition to MK-D1 ESCRT II recruitment at these regions, MK-D1 ESCRT-III and VPS4 form ring assemblies at midbodies and target reforming nuclear envelope regions. Futhermore, expression of MK-D1 ESCRT modules enhances efficiency of viral budding in human cells. These findings provide direct experimental evidence that archaeal ESCRT modules integrate into diverse eukaryotic structures, supporting a model in which ancestral membrane-remodeling systems were redeployed during eukaryogenesis to organize novel internal architectures.

## Introduction

The origin of eukaryotic cellular complexity remains a central question in evolutionary biology. A defining feature of eukaryotic cells is the presence of internal membrane systems, including the endoplasmic reticulum, endosomes, and the nuclear envelope, which are absent from bacteria and archaea. The molecular mechanisms underlying the emergence of these compartments during eukaryogenesis remain largely unresolved, particularly how ancestral proteins were adapted or repurposed for novel roles within the expanded architectural complexity. The discovery of Asgard archaea and later and cultivation of *Prometheoarchaeum syntrophicum* (MK-D1), and later again *Candidatus Lokiarchaeum ossiferum, Candidatus Margulisarchaeum peptidophila* and *Candidatus Flexarchaeum multiprotrusionis* has yielded critical insight into the archaeal-to-eukaryotic transition^1–5^. These organisms are morphologically simple, lacking internal compartments and bounded by a single cell membrane, yet their genomes encode numerous eukaryotic signature proteins (ESPs), including those implicated in cytoskeletal dynamics and membrane remodeling. Despite extensive genomic and structural insights, how simple archaeal cells evolved into the compartmentalized architecture of eukaryotes remains one of the most persistent mysteries in biology. This raises a fundamental question: how were ancestral Asgard archaeal proteins co-opted for roles in the development of internal membrane systems?

Among the many systems that remodel membranes, the ESCRT machinery stands out for its evolutionary conservation and mechanistic versatility. Originating early in the archaeal lineage^6^, ESCRT complexes in eukaryotes have diversified to orchestrate abscission, endosomal sorting, and nuclear envelope remodeling core processes that define the internal architecture of complex cells. In eukaryotes, the ESCRT (Endosomal Sorting Complex Required for Transport) machinery mediates membrane remodeling and scission across diverse cellular processes, including cytokinetic abscission, endosomal cargo sorting, nuclear envelope reformation, and viral budding^7–9^. The ESCRT-II complex, composed of VPS22, VPS25, and VPS36, stabilizes membrane curvature and acts as a recruitment platform for ESCRT-III subunits. VPS36 also binds ubiquitinated cargo, facilitating membrane sorting. ESCRT-III components (CHMP1–CHMP7) assemble into dynamic polymers at membrane necks, with CHMP4/SNF7 forming the core filament that drives membrane constriction and fission^10^. The AAA+ ATPase VPS4 catalyzes the disassembly and recycling of ESCRT-III polymers following scission. ESCRT-III assembly during cytokinesis is coordinated by ALIX and ESCRT-I/II, which are recruited to the midbody by CEP55^11–13^.

However, whether archaeal ESCRT homologues retain functional or localization compatibility with eukaryotic membrane systems has remained untested. To bridge the gap between sequence homology and functional capacity, we experimentally tested whether ESCRT proteins from *P. syntrophicum* (MK-D1) could operate within a eukaryotic cellular context. It remains unknown whether their proteins are intrinsically capable of functioning within the diverse internal architectures of eukaryotic cells. To investigate the evolutionary continuity and possible functional compatibility of archaeal ESCRT proteins in a eukaryotic context, we heterologously expressed fluorescently tagged MK-D1 ESCRT proteins in human HeLa cells and assessed their subcellular localization. Specifically, we asked whether these archaeal proteins could recognize and be recruited to internal structures and remodeling sites characteristic of eukaryotic cells, despite the absence of these features in MK-D1 cells. In addition to structural similarity, we show that ESCRT-II, ESCRT-III, and VPS4 proteins from MK-D1 localize to centrosomes, midbodies, nuclear envelopes, endosomes, and viral budding sites in human cells. These findings provide direct evidence that archaeal ESCRT proteins are intrinsically compatible with eukaryotic membrane systems and can be redeployed to multiple eukaryotic sites, supporting a model in which ancestral membrane-remodeling systems were repurposed during eukaryogenesis to organize novel internal architectures.

## Results

### MK-D1 Encodes Ubiquitin and ESCRT-II Subunits with Conserved Cores and Divergent Regulatory Domains

MK-D1 protein coding sequences for ubiquitin (UB), ESCRT-II subunits (VPS22, VPS25, VPS36)^10,14^, were identified by BLAST searches and confirmed to possess structural homology to their human counterparts, through phylogenetic analysis, structure prediction in AlphaFold3 (AF3) or by X-ray crystallography in the case of VPS25 (Figures 1 and S1; Tables S1 and S2)^1,15^. The X-ray structure of VPS25 closely reproduced the structures of Odinarchaeota, yeast and human VPS25s, with differences in the relative angles of the two winged-helix domains (WH D1 and D2, Figure 1B), while the MK-D1 VPS25 AF3 model indicates an unstructured N-terminal extension. The MK-D1 VPS36 AF3 model also reveals a shared ancestral structural core comprised of two WH domains (Figure 1C). However, divergence in their N-terminal domains, MK-D1 VPS36 harbors an oligosaccharide-binding fold (OBF) whereas hVPS36 has a ubiquitin-binding domain, suggests that cargo recognition mechanisms and regulatory features have evolved independently. Similarly, both the MK-D1 AF3 model and human VPS22 (hVPS22) structure contain the core WH domains but are preceded by divergent N-terminal helical motifs (Figure 1D). The predicted MK-D1 ubiquitin fold is broadly conserved with differences in loop regions (Figure 1E). Taken together, MK-D1 encodes ubiquitin and ESCRT-II subunits with conserved cores but divergent N-terminal domains. These observations suggest that the structural framework of eukaryotic ESCRT-II and ubiquitin systems were already established in Asgard archaea, while regulatory and cargo-recognition features diversified later during eukaryogenesis.

**Figure 1.**
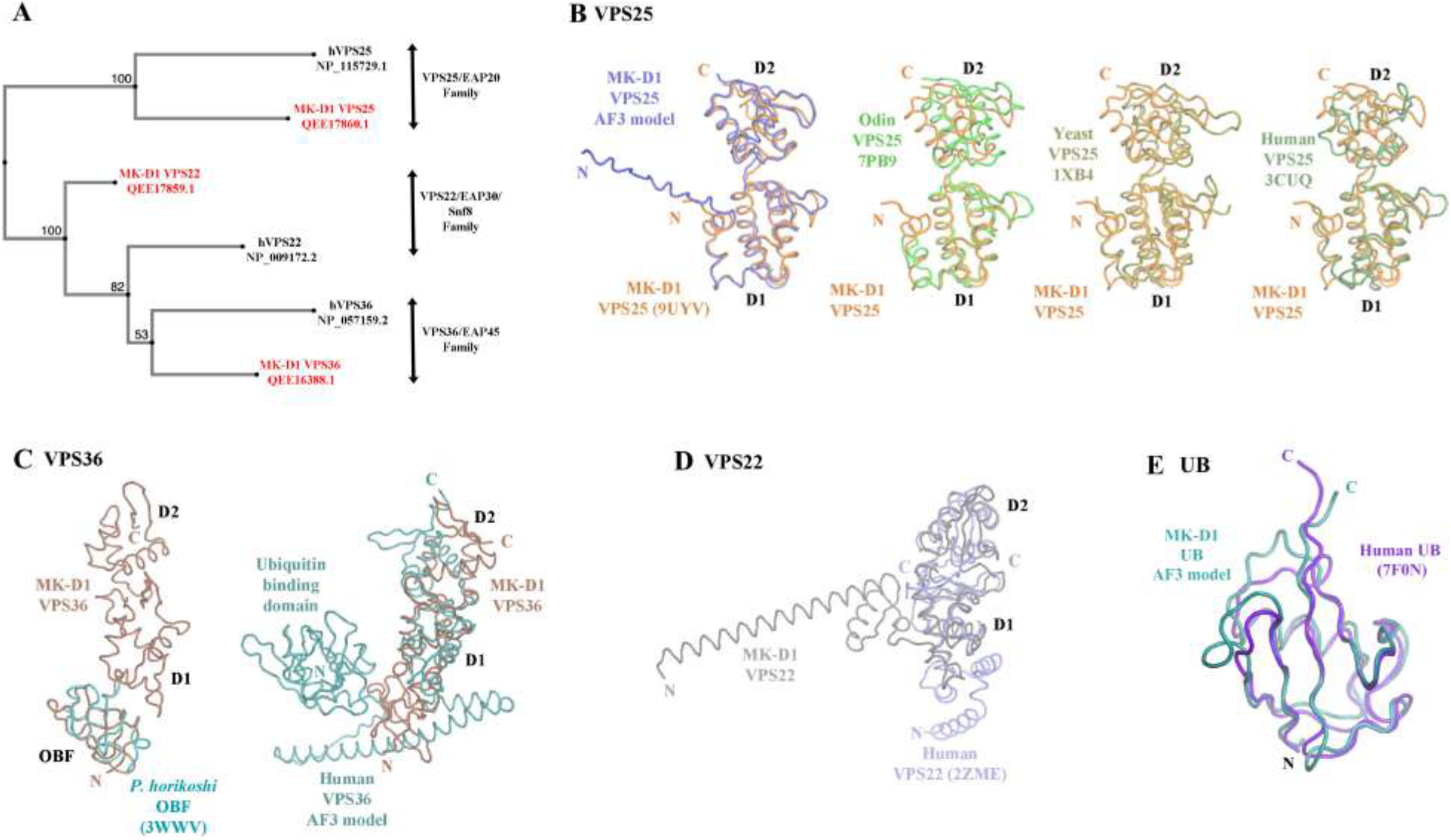
Phylogenetic and structural comparisons of MK-D1 ESCRT-II proteins and UB with human homologs. (A) Phylogenetic analysis of the MK-D1 ESCRT II proteins with human counterparts. MK-D1 ESCRT-II were named in this study based on their nearest phylogenetic neighbors. Structural comparisons of MK-D1 ESCRT-II and ubiquitin proteins predicted by AF3. (B–E) Structural comparisons of MK-D1 ESCRT II proteins with human homologs. (B) The X-ray structure of MK-D1 VPS25 closely matches the AF3 prediction, though the model suggests an extended N-terminus. MK-D1 VPS25 also superimposes well with VPS25 structures from Odinarchaeota, yeast, and human, with variable angles between domain 1 (D1) and domain 2 (D2). (C) AF3-predicted MK-D1 VPS36 contains an N-terminal oligosaccharide binding fold (OBF), and D1 and D2 superimpose with these domains from human VPS36, although the angles D1 between D2 differ. However, human VPS36 contains an N-terminal ubiquitin-binding domain and an additional helical segment. (D) The AF3-predicted MK-D1 VPS22 D1 and D2 superimpose well with the human counterparts, although differences are observed in the N-terminal helices. (E) The AF3-predicted MK-D1 ubiquitin predicted structure closely matches the human ubiquitin structure, with differences in loop regions.

### MK-D1 ESCRT-II homologs integrate into centrosomal, midbody, and endosomal structures

To assess the functional implications of this evolutionary divergence, we next asked whether MK-D1 ESCRT-II components can operate in the eukaryotic cellular environment. Specifically, we investigated their localization and behavior in human cells. First, we examined the localization of fluorescently tagged MK-D1 VPS36 in HeLa cells. We consistently observed an intense ∼1-2 μm fluorescence spot for MK-D1 VPS36 in the cytoplasm, which colocalized with a marker for the centrosome, human MKKS centrosome shuttling protein, and with human ESCRT-II VPS25 (hVPS25) (Figures 2A and S2A)^16^. Similar intense MK-D1 VPS36 and hVPS25 foci colocalized with a second marker for centrosomes, centrosomal protein 55 (CEP55) (Figures 2B and S2B), which were sometimes observed as a pair of fluorescent foci, likely due to cell cycle-dependent centrosome duplication (Figure 2C). This pattern was consistently observed across independent experiments, arguing against random aggregation. CEP55 is implicated in centrosome-dependent cellular processes, such as centrosome duplication, cell cycle progression, and cytokinesis regulation^17^. Other cells showed greater numbers of MK-D1 VPS36 foci, which colocalized with a marker for centriolar satellites, centrosomal protein 131 (CEP131), and with hVPS25 (Figure 2D). Similarly, MK-D1 and human VPS22 (hVPS22) ESCRT-II proteins colocalized to centrosome structures (Figures S2C and S2D). These data demonstrate that human and MK-D1 ESCRT-II components are recruited to centrosomal (CEP55) and pericentrosomal (CEP131) regions when expressed in HeLa cells. The coordinated localization of multiple MK-D1 ESCRT-II subunits with distinct centrosomal markers and their cell cycle–dependent duplication patterns collectively demonstrate that this recruitment is specific and organized suggesting association is not a passive consequence of overexpression but reflects conserved molecular recognition of centrosomal organizing cues. This unexpected recruitment of ESCRT-II proteins to centrosomes suggests that centrosomal ESCRT activity may be more widespread than currently appreciated.

**Figure 2.**
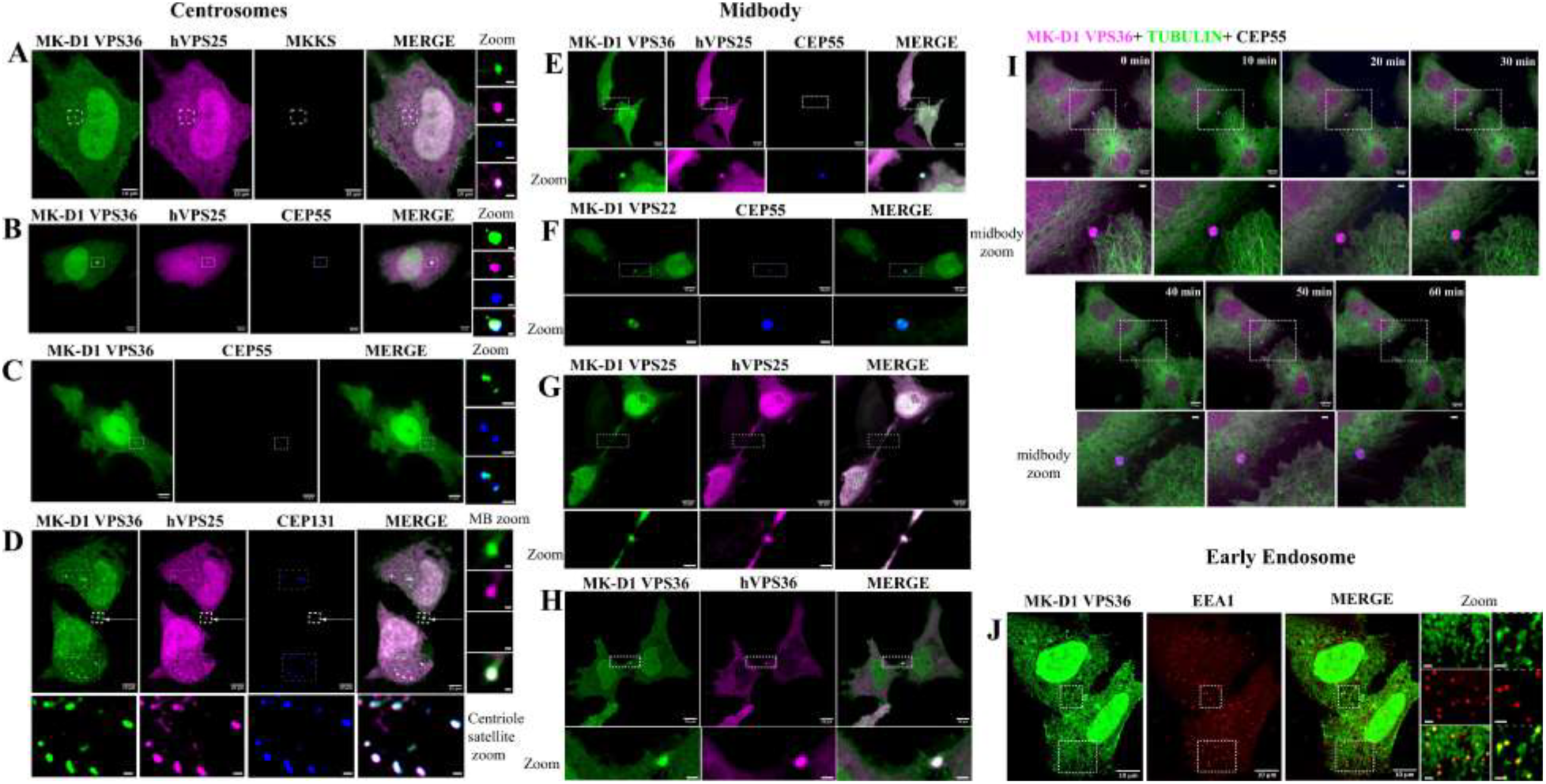
MK-D1 ESCRT-II homologs integrate into centrosomal, midbody, and endosomal structures. Representative images of HeLa cells co-transfected with fluorescently tagged MK-D1 and human ESCRT-II constructs, followed by confocal and super-resolution microscopy. Centrosome: (A) EGFP-MK-D1 VPS36, mCherry-human VPS25 (hVPS25) and the centrosomal shuttling protein TagBFP-MKKS; (B) EGFP-MK-D1 VPS36, mCherry-hVPS25 and centrosomal marker TagBFP-CEP55; (C) EGFP-MK-D1 VPS36 and TagBFP-CEP55, showing cell cycle–dependent centrosome duplication; (D) EGFP-MK-D1 VPS36, mCherry-hVPS25, and the centriolar satellite marker TagBFP-CEP131 localizing to multiple foci. Midbody (MB): (E) EGFP-MK-D1 VPS36, mCherry-hVPS25, and TagBFP-CEP55 colocalize at structures separated from the main cell body; (F) Asymmetric colocalization of EGFP-MK-D1 VPS22 with TagBFP-CEP55, suggesting midbody substructure; (G) EGFP-MK-D1 VPS25 and mCherry-hVPS25 enriched at the midbody. (H) EGFP-MK-D1 VPS36 colocalizing with mCherry-hVPS36 at the midbody. (I) Time-lapse super-resolution imaging of mCherry-MK-D1 VPS36, EGFP-tubulin and TagBFP-CEP55, showing midbody inheritance and internalization into a cell (Movie S1). (J) EGFP-MK-D1 VPS36 colocalization with the early endosomal marker mCherry-EEA1, indicating recruitment to early sorting vesicles. Scale bars: main images 10 μm, zoom images 2 μm.

We often observed a ∼2 μm structure separated from the cell body that also showed high intensity for fluorescently tagged MK-D1 VPS36 and hVPS25, which was not positive for CEP131 (Figure 2D). CEP55, which is present at centrosomes during interphase and early mitosis but moves to the midbody as mitosis concludes^18^, co-localized with MK-D1 VPS36, human VPS36 (hVPS36), hVPS25 and MK-D1 VPS25 in these peripheral structures confirming their identity as the midbody (Figures 2E, S2E and S2F). MK-D1 VPS22 and hVPS22 also colocalized with the midbody markers CEP55 and MK-D1 VPS36, respectively (Figures 2F, S2C, S2D and S2G), with one image showing duplet fluorescence foci for MK-D1 VPS22 that were symmetrically separated to two poles of the CEP55-marked midbody, indicating possible substructure (Figure 2F). Expression of pairs of VPS22, VPS25 and VPS36 proteins confirmed that all components of the human and MK-D1 ESCRT-II complexes were enriched at the midbody (Figures 2E-2H, S2C, S2D and S2F-S2I). Unlike the case for the centrosome, the presence of human ESCRT-II proteins at the midbody is well established^19^.

We then examined whether MK-D1 VPS36-associated midbodies were associated with microtubules. MK-D1 VPS36-labeled midbodies showed overlap, or partial overlap, with tubulin at the midbody (Figures S3A and S3B). As the MK-D1 VPS36-positive midbody often appeared asymmetric and in close proximity to one of the daughter cells (Figures 2H and S3C), we followed the fate of the midbody by tracking MK-D1 VPS36 localization. Over a period of approximately 1 hour, a midbody was observed to be asymmetrically internalized (Figure 2I; Movie S1), consistent with previous findings that postmitotic midbodies can be inherited and internalized^20^. Furthermore, the MK-D1 VPS36 pattern at the midbody became more asymmetric over time indicating dynamic remodeling of midbody substructures (Figure 2I).

In addition, MK-D1 ESCRT II VPS36 colocalized with the early endosomal marker EEA1, showing recruitment to early sorting vesicles (Figure 2J), consistent with roles linking ESCRT assemblies to membrane trafficking. Together, these results indicate that MK-D1 ESCRT-II homologs are structurally conserved and localized to centrosomes, midbodies, and endosomes in human cells suggesting that these proteins had evolved essential properties prior to the emergence of eukaryotes. These observations demonstrate that MK-D1 ESCRT-II components are not passively distributed but instead respond to the eukaryotic cellular context, and that their asymmetric localization is consistent with functional recruitment rather than random aggregation. Taken together, MK-D1 ESCRT-II homologs function as adaptive modules across diverse subcellular pathways, suggesting ancestral framework for eukaryotic membrane remodeling and protein trafficking.

### MK-D1 ESCRT-III and VPS4 homologs

MK-D1 protein coding sequences for ESCRT-III subunits (CHMP4, CHMP1A, CHMP1B, VPS4) were identified by BLAST searches and confirmed to possess structural homology to their human counterparts, through phylogenetic analysis and structure prediction in AlphaFold3 (AF3)^15^ (Figures 3 and S1). The MK-D1 ESCRT-III subunits are predicted to possess the canonical helix-turn-helix motif (Figures 3B and 3C). VPS4 is predicted to closely reproduce the eukaryotic homolog structures, albeit the relative orientations of the VPS4 domains are predicted to be different (Figure 3D). Taken together, this phylogenetic and structure analysis suggests that while the ESCRT III and VPS4 systems from MK-D1 and eukaryotes share a common origin, they exhibit potential structural variations resulting from lineage diversification of the membrane-remodeling machinery (Figure 3).

**Figure 3.**
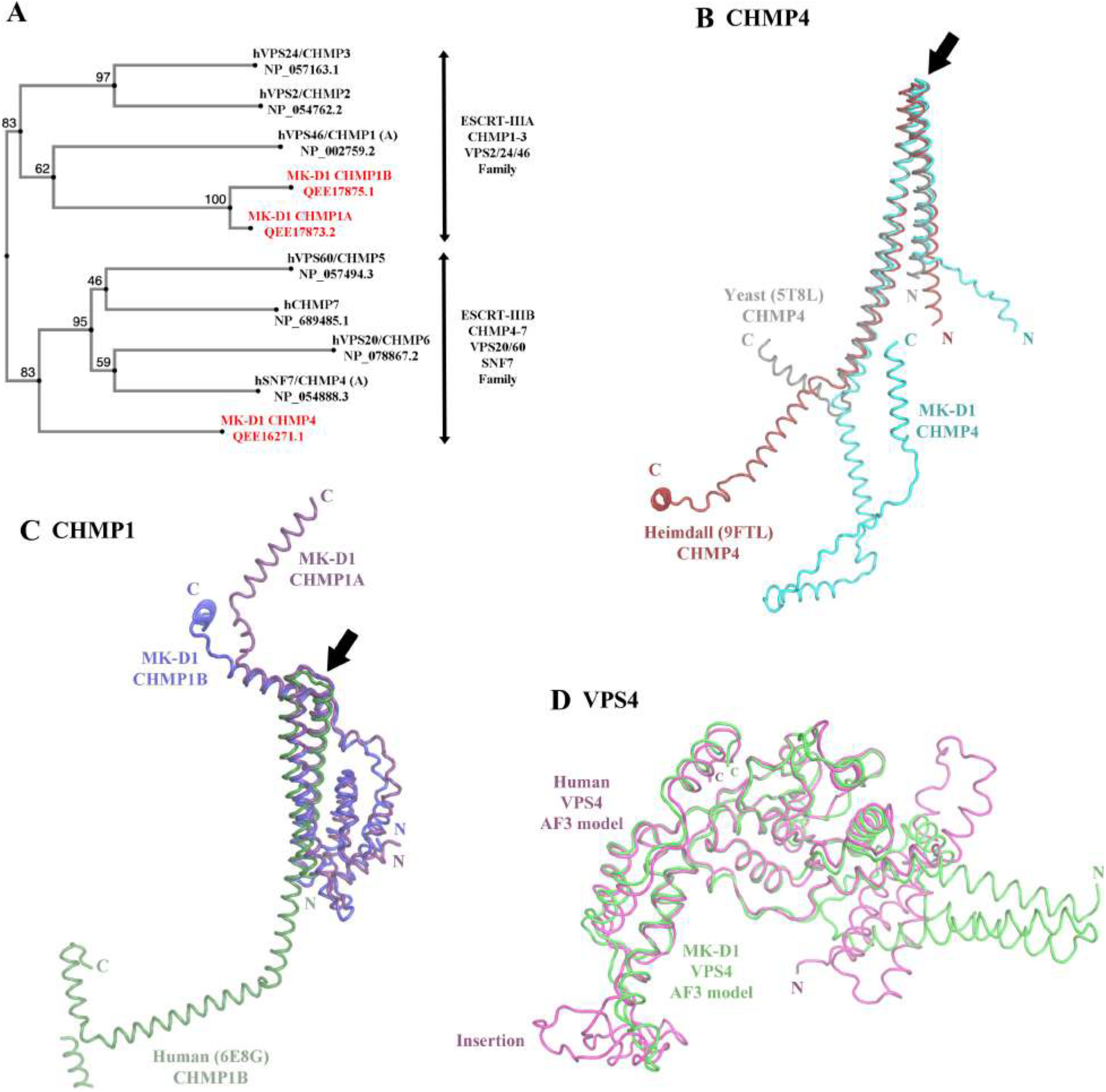
Phylogenetic and structural comparisons of MK-D1 ESCRT-III proteins with human homologs. (A) Phylogenetic analysis of the MK-D1 ESCRT III proteins with human counterparts. MK-D1 ESCRT-III were named in this study based on their nearest phylogenetic neighbors. (B–E) Structural comparisons of MK-D1 ESCRT III protein AF3-predicted structures with human homologs. (B-C) AF3-predicted ESCRT-III proteins show good alignment of the helix-turn-helix motifs (arrows), while the remaining regions are flexible and likely dependent on complex formation. (D) AF3-predicted MK-D1 and human VPS4 structures superimpose well in their ATPase domains. hVPS4 has an insertion not present in MK-D1, and the predicted N-terminal helical domain orientations differ.

### MK-D1 ESCRT-III and VPS4 homologs integrate into centrosomal, midbody, and endosomal structures

We next examined the localization of MK-D1 ESCRT-III proteins and the ATPase VPS4 in relation to centrosomes and midbodies, sites where the recruitment of their human homologs has been previously demonstrated^10,13,19,21–23^. In HeLa cells expressing fluorescently tagged MK-D1 CHMP4, we observed co-localization with the centrosomal markers CEP55, hVPS4, and spastin (Figures 4A and 4B). Midbody co-localization was observed for MK-D1 CHMP4 with hVPS4 and CEP55, MK-D1 VPS4 with hVPS4 and the human ESCRT-III component CHMP3 (hCHMP3), and MK-D1 CHMP1A with hVPS4 (Figures 4C-4F). Given that CEP55 is a shared component of both the centrosome and midbody, it likely facilitates the recruitment of ESCRT-II/III and VPS4 through interactions with ESCRT-I and ALIX in both structures, as has been described in cytokinetic abscission^12^. In addition, MK-D1 VPS4 colocalized extensively with the early endosomal marker EEA1 (Figure 4G), compared to ESCRT II (Figure 2J), showing recruitment to early sorting vesicles and suggesting that MK-D1 VPS4 preferentially functions in early endosomal sorting, suggesting an ancestral, simplified ESCRT deployment. Taken together, these findings indicate that MK-D1 ESCRT-III proteins and VPS4 are not only structurally conserved with their eukaryotic counterparts but also dynamically recruited to centrosomes, midbodies, and endosomal compartments, supporting modular deployment model, originating prior to eukaryogenesis, in which ESCRT-II provides organizational stability, while ESCRT-III and VPS4 contribute dynamic functions at diverse membrane remodeling sites.

**Figure 4.**
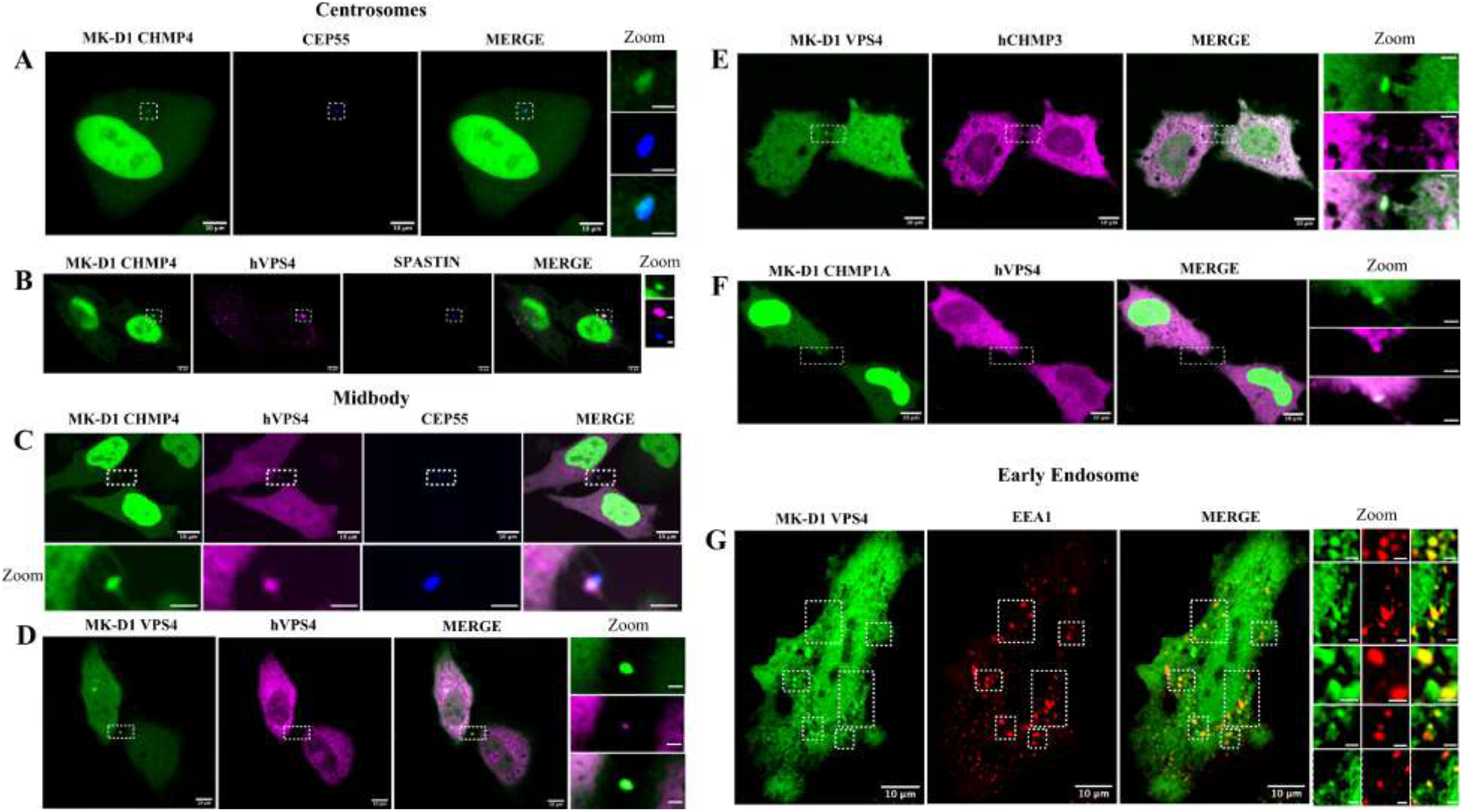
Representative images of HeLa cells co-transfected with fluorescently tagged MK-D1 and human ESCRT-III constructs, followed by confocal and super-resolution microscopy. Centrosome: (A) EGFP-MK-D1 CHMP4 and centrosomal marker TagBFP-CEP55; (B) EGFP-MK-D1 CHMP4, mCherry-hVPS4 and TagBFP-spastin; Midbody: (C) EGFP-MK-D1 CHMP4 and mCherry-hVPS4 colocalize with TagBFP-CEP55; (D) EGFP-MK-D1 VPS4 and mCherry-hVPS4; (E) EGFP-MK-D1 VPS4 and mCherry-hCHMP3. (F) EGFP-MK-D1 CHMP1A and mCherry-hVPS4. (G) EGFP-MK-D1 VPS4 localization with the early endosomal marker mCherry-EEA1 indicating recruitment to early sorting vesicles. Scale bars: main images 10 μm, zoom images 2 μm.

### MK-D1 ESCRT-III/VPS4 recapitulate midbody ring-like structures localize to nuclear envelopes

MK-D1 ESCRT-III/VPS4 homologs closely resemble their human counterparts and localize to centrosomes, midbodies, and endosomal compartments (Figures 3 and 4). However, it is not yet clear whether they can assemble functional midbody rings, engage the abscission machinery, or contribute to nuclear envelope remodeling in eukaryotic cells. To address this, we investigated their roles in ESCRT-mediated cytokinesis and nuclear remodeling. High-resolution imaging of fluorescently tagged MK-D1 CHMP4 revealed ∼2 μm diameter curved structures at the midbody (Figures 5A and 5B), consistent with its predicted helix-turn-helix polymerization motif (Figure 4B). This spatial arrangement aligns with native ESCRT-III activity in late mitosis, wherein polymerizing CHMP proteins form < 2 μm diameter constriction rings that mediate daughter cell separation^24,25^. The MK-D1 CHMP4 partial ring structures co-localized with MK-D1 VPS4 and tubulin (Figure 5A; blue arrow), and with CEP55 and MK-D1 VPS4 (Figure 5B; blue arrow). These patterns are reminiscent of ESCRT-III deployment during eukaryotic cytokinesis and suggest that MK-D1 ESCRT-III homologs may retain functional features involved in membrane fission^12,25,26^. We explored whether other foci nearby MK-D1 CHMP4 partial ring structures delineate the abscission zone, raising the possibility that this protein contributes to membrane fission. In one dividing cell pair, a patch of MK-D1 CHMP4 lacking VPS4 was observed adjacent to the main ring (Figure 5B; orange arrow), whereas in another pair, MK-D1 CHMP4 co-localized with VPS4 in this region (Figure 5A; orange arrow), suggesting possible temporal recruitment of VPS4 to these sites. The diameters of potential abscission sites were less than a half of the diameter of the main MK-D1 CHMP4 rings, consistent with the size of abscission site rings observed in HeLa cells^24^. Although we were unable to capture the extension of MK-D1 CHMP4 assemblies from the midbody to the abscission zone in real time, likely due to the narrow temporal window of this event, MK-D1 ESCRT-II/III and VPS4 consistently localized to the midbody and formed distinct substructures reminiscent of eukaryotic ESCRT activity^22,25^. Together, these observations support a modular deployment of the MK-D1 ESCRT-III and VPS4 contributing to dynamic functions related to membrane scission.

**Figure 5.**
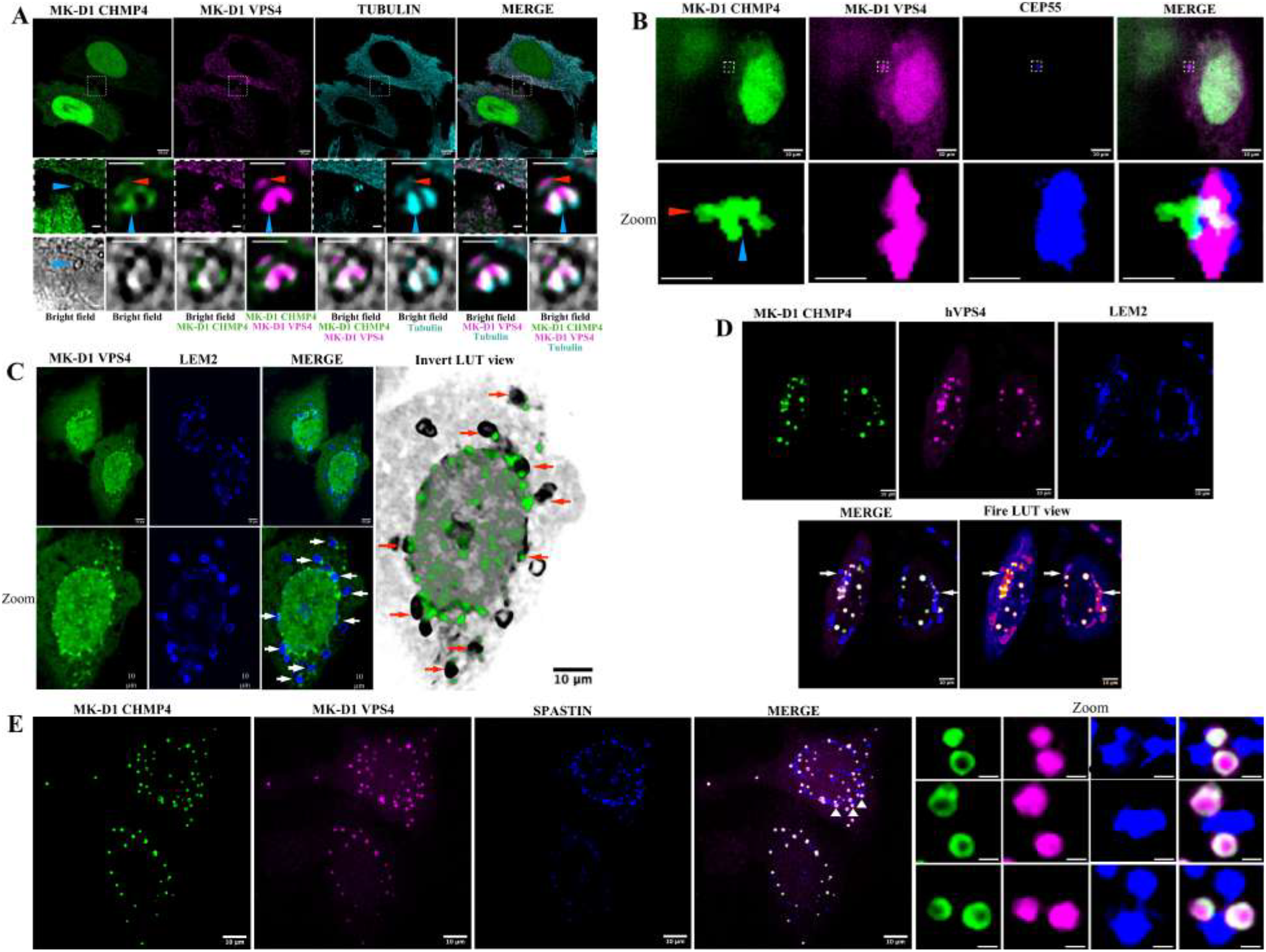
MK-D1 ESCRT-III/VPS4 recapitulate midbody ring assembly, associate with abscission zones, and localize to the reforming nuclear envelope. Representative images of HeLa cells co-transfected with fluorescently tagged MK-D1 and human ESCRT-III or VPS4 constructs, followed by confocal and super-resolution microscopy. (A-B) Super-resolution imaging reveals ring-like assemblies of MK-D1 CHMP4 in the midbody region (blue arrowheads) and potential abscission zone (orange arrowheads). (A) EGFP-MK-D1 CHMP4, mCherry-MK-D1 VPS4, and SPY650 tubulin. (B) EGFP-MK-D1 CHMP4, mCherry-MK-D1 VPS4, and TagBFP-CEP55. Orange arrowheads in a and b indicate potential abscission zones adjacent to the CHMP4 ring. Scale bars: main images 10 μm, zoom images 2 μm. (C-E) MK-D1 ESCRT-III proteins and VPS4 localize to the reforming nuclear envelope during mitotic exit. (C) EGFP-MK-D1 VPS4 forms punctate foci at the reforming nuclear envelope in dividing cells, co-localizing with or adjacent to the inner nuclear membrane marker TagBFP-LEM2 (arrows). The inverted Lookup Table (LUT) view enhances contrast at membrane boundaries, facilitating their delineation. (D) EGFP-MK-D1 CHMP4 co-localizes with mCherry-hVPS4 near TagBFP-LEM2 foci at the nuclear periphery (example sites, arrows). (E) Super-resolution imaging reveals circular foci composed of EGFP-MK-D1 CHMP4 and mCherry-MK-D1 VPS4 form at the nuclear envelope during mitotic exit, often adjacent to or overlapping with the microtubule-severing enzyme TagBFPastin. Insets show high-magnification views of candidate droplets of MK-D1 CHMP4 and MK-D1 VPS4 adjacent to spastin foci (arrowheads). Scale bars: main images 10 μm, zoom scale bar: 1 μm.

We next examined whether MK-D1 ESCRT homologs are recruited to the reforming nuclear envelope during mitotic exit. In eukaryotic cells, ESCRT-III proteins are recruited to the reforming nuclear envelope, where they mediate membrane sealing and reestablish nuclear compartmentalization^27,28^. This process is coordinated by the inner nuclear membrane protein LEM2, which recruits CHMP7, an ER-localized ESCRT-II/ESCRT-III hybrid protein^29,30^, and the microtubule-severing enzyme spastin, which together facilitate the recruitment and activity of ESCRT-III components. In dividing HeLa cells, MK-D1 VPS4 formed discrete foci at the reforming nuclear envelope, frequently overlapping or adjacent to LEM2-positive sites (Figures 5C, S4A and S4B). MK-D1 CHMP4 also co-localized with hVPS4 near LEM2 clusters (Figures 5D and S4C). In addition, peripheral nuclear foci containing both MK-D1 CHMP4 and VPS4 were frequently observed adjacent to spastin foci (Figure 5E). A similar spatial relationship was observed for MK-D1 CHMP4, hVPS4, and spastin (Figures S4D and S4E). These observations indicate that archaeal ESCRT components can recapitulate key spatial interactions involved in nuclear envelope remodeling, supporting the hypothesis that the ancestral ESCRT-III machinery was co-opted during eukaryotic evolution for this specialized membrane remodeling function. At the nuclear periphery, MK-D1 CHMP4/VPS4 foci appeared spherical, possibly indicating liquid–liquid phase separation (Figures 5E, S5A and S5B), a hallmark of native LEM2/ESCRT-III complexes involved in nuclear envelope sealing^31^. Supporting this, PICNIC-based predictions yielded similar condensate-localization probabilities for MK-D1 and human CHMP4 (0.70 vs. 0.81) and VPS4 (0.77 vs. 0.80), while human LEM2 (0.24) scored below the condensation threshold (0.5)^32^. These results suggest MK-D1 CHMP4 and VPS4 exhibit the biophysical behaviors of their eukaryotic counterparts. Taken together, our results demonstrate that MK-D1 ESCRT-III proteins and VPS4 are structurally conserved, dynamically recruited to key membrane sites, and capable of recapitulating eukaryotic-like spatial interactions and biophysical behaviors. This supports a modular deployment model and suggests that the ancestral ESCRT-III machinery was co-opted during eukaryotic evolution to mediate specialized processes such as nuclear envelope sealing and membrane scission.

### MK-D1 UB–ESCRT modules display compartment specific redeployment across cellular and viral pathways

We asked how MK-D1 ubiquitin and ESCRT modules achieve compartment-specific recruitment in human cells, and what this reveals about the evolutionary origins of membrane-remodeling pathways. Given that the UB–ESCRT pathway is a well-established mechanism in eukaryotic cells, we specifically examined the localization and coordination of ubiquitin-associated ESCRT-II, ESCRT-III, and VPS4 at midbodies, centrosomes, the nuclear envelope, endosomes, and viral release sites to understand their context-dependent deployment. MK-D1 VPS36 localized to the CEP55-marked midbody (Figure 6A) and fluorescently tagged MK-D1 UB consistently co-localized with MK-D1 VPS36 and MK-D1 VPS22 at the midbody (Figures 6B and 6C). MK-D1 UB was also recruited to centrosomes (Figure 6B) and centriolar satellites marked by CEP131 (Figure S6A). Tubulin was also present at centriolar satellites (Figure S6B), suggesting a potential role of UB in organizing centrosomal architecture to support microtubule dynamics. MK-D1 ESCRT-II component VPS36 co-localized with ESCRT-III hCHMP4 and tubulin at the midbody, forming rings and patches at the potential abscission zone (Figure S6C). Similarly, MK-D1 CHMP4 co-localized with tubulin at the midbody in rings and patches (Figure 5A) and was also observed together with MK-D1 VPS4 (Figure 5A) or positioned adjacent to it (Figure 5B), consistent with a model of sequential deployment. Together, these results suggest that MK-D1 UB can serve as a spatial molecular cue to recruit ESCRT-II components during midbody-associated membrane remodeling.

**Figure 6.**
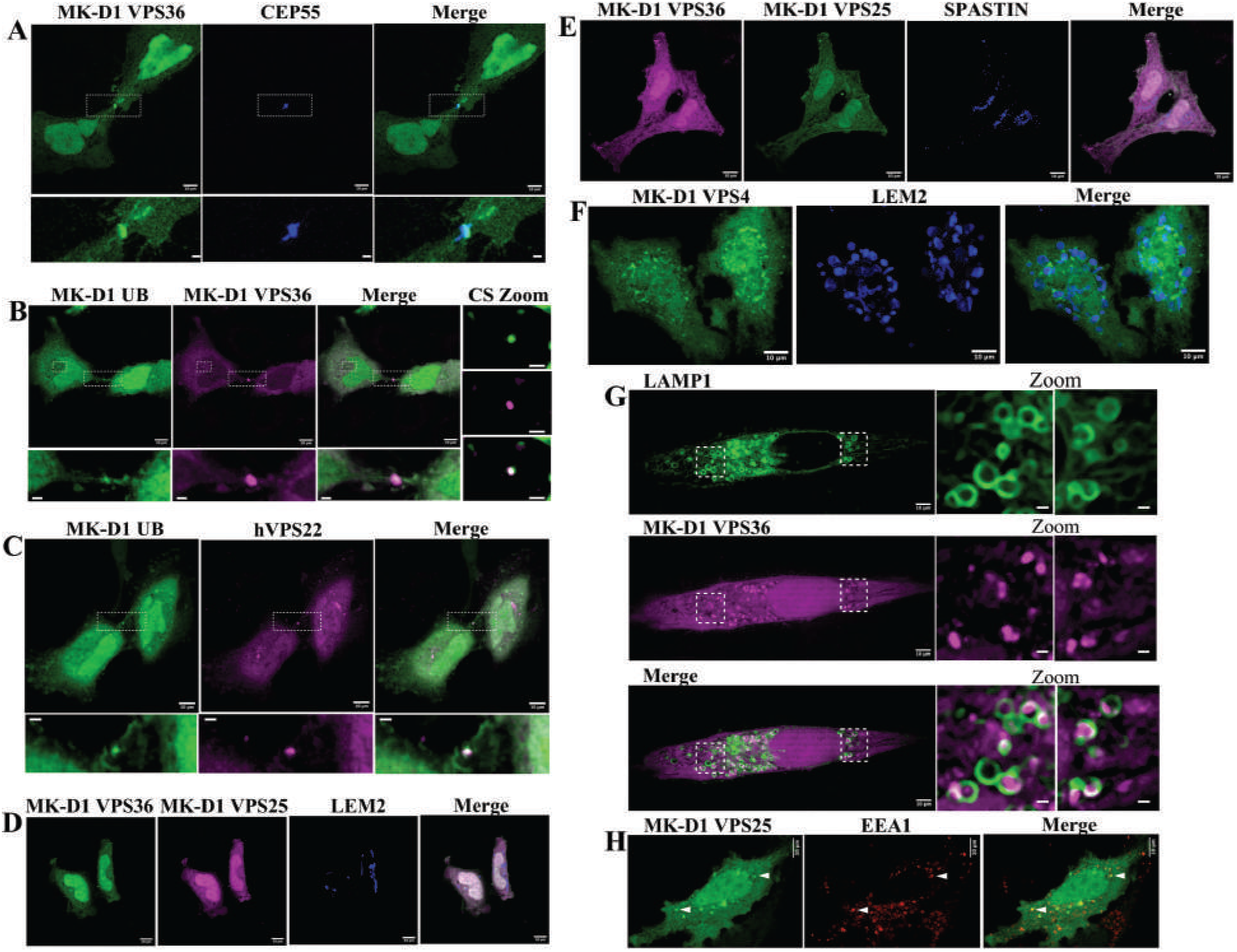
Compartment-specific deployment of MK-D1 UB–ESCRT modules across cellular and viral contexts. Representative images of HeLa cells co-transfected with fluorescently tagged MK-D1 UB and ESCRT constructs, followed by confocal and super-resolution microscopy. (A) EGFP-MK-D1 VPS36 localizes with TagBFP-CEP55 at the midbody. (B) EGFP-MK-D1 UB and mCherry-hVPS36 at the midbody. (C) EGFP-MK-D1 UB localizes with mCherry-hVPS22 at midbody. (D) EGFP-MK-D1 VPS36 and mCherry-MK-D1 VPS25 do not localizes with the inner nuclear membrane marker TagBFP-LEM2, or with (E) spastin, indicating exclusion of ESCRT-II from nuclear envelope remodeling. (F) EGFP-MK-D1 VPS4 localizes with the inner nuclear membrane marker TagBFP-LEM2. (G) EGFP-MK-D1 VPS36 localizes to vesicles positive for mCherry-LAMP1A (magenta), a marker of late endosomes and lysosomes. (H) EGFP-MK-D1 VPS25 shows partial colocalization with the early endosomal marker mCherry-EEA1.

At reforming nuclear envelope regions, MK-D1 ESCRT-II proteins VPS25 and VPS36 did not form puncta near LEM2 or spastin foci (Figures 6D and 6E), indicating that MK-D1 ESCRT-II is not engaged at these sites, in contrast to the site-specific recruitment observed for MK-D1 ESCRT-III and VPS4 supporting a compartment-specific deployment model (Figures 5C–5E and 6F). In eukaryotic cells, the ESCRT-II VPS36 interaction with UB is the initial signaling cue in sorting ubiquitinated cargo at early endosomes and promoting multivesicular body (MVB) formation, ultimately linking the ESCRT pathway to lysosomal degradation^33–37^. Despite the structural similarity of MK-D1 UB with human UB (Figure 1E), MK-D1 ubiquitin was not detected at endosomes likely reflecting divergence in VPS36 N-terminal domains, as MK-D1 VPS36 carries an OBF fold rather than the ubiquitin-binding domain of hVPS36 (Fig 1 C and E), suggesting that cargo recognition and regulation evolved independently.

Nevertheless, MK-D1 ESCRT-II VPS36, which did not form puncta at the reforming nuclear envelope, was found to co-localize with LAMP1A-positive lysosomes, similar to hVPS36 (Figures 6G, S6D and S6E), and MK-D1 ESCRT-II proteins VPS25 and VPS36 localized to some EEA1-positive early endosomes (Figures 6H and 2J). Additionally, hVPS4 and MK-D1 VPS4, which forms discrete puncta, also co-localized with endosomes (Figure 4G) and lysosomes (Figures S6F and S6G). Thus, MK-D1 ESCRT-II is excluded from nuclear envelope remodeling but recruited to endo-lysosomal compartments, highlighting a compartment-specific, modular deployment of ESCRT machinery.

In eukaryotic cells, viral budding involves ubiquitination of viral proteins and recognition by ESCRT-II^38,39^. We therefore asked if MK-D1 ESCRT homologs could also localize to viral assembly sites. To test this, mCherry-tagged MK-D1 VPS25 and VPS4 were expressed in retrovirus-producing Plat-GP cells together with VSV-G and a retroviral genome, and their localization was examined relative to the viral structural protein Gag. Since Gag interacts with ESCRT-I and ALIX and mimics the ESCRT-0–ESCRT-I interaction^40^, we reasoned that it could potentially recruit archaeal ESCRT homologs. Indeed, both mCherry-tagged MK-D1 VPS25 and VPS4 displayed increased co-localization with Gag compared to mCherry alone, suggesting targeted recruitment to viral assembly sites (Figure S7). To assess functional relevance, we measured viral output in infection assays. Expression of MK-D1 VPS25 and VPS36, but not VPS4 or CHMP4, resulted in a significant increase in viral release, indicating that these ESCRT-II homologs may contribute to the budding process (Figure 7A). Structural analyses indicate that MK-D1 VPS36 lacks the canonical ubiquitin-binding domain and instead contains an OBF fold (Figure 1C), suggesting that viral recruitment may not rely on direct ubiquitin recognition. This suggests that viral recruitment evolved independently, but VPS25 and VPS36 retain sufficient modularity to function in this context, this is consistent with their previously observed localization to endosomes, and points to a conserved deployment of VPS25 and VPS36 across both viral and endolysosomal pathways. Our findings reveal that MK-D1 ESCRT modules are flexibly deployed across centrosomal, midbody, endosomal, nuclear, and viral pathways. Taken together, this study suggests that mechanistic deployment model (Figure 7B), in which ancestral UB–ESCRT proteins were redeployed across distinct cellular contexts. Such compartment-specific redeployment highlights an ancestral modular system that predates eukaryotic complexity and provides insight into the evolutionary trajectory of membrane-remodeling machinery.

**Figure 7.**
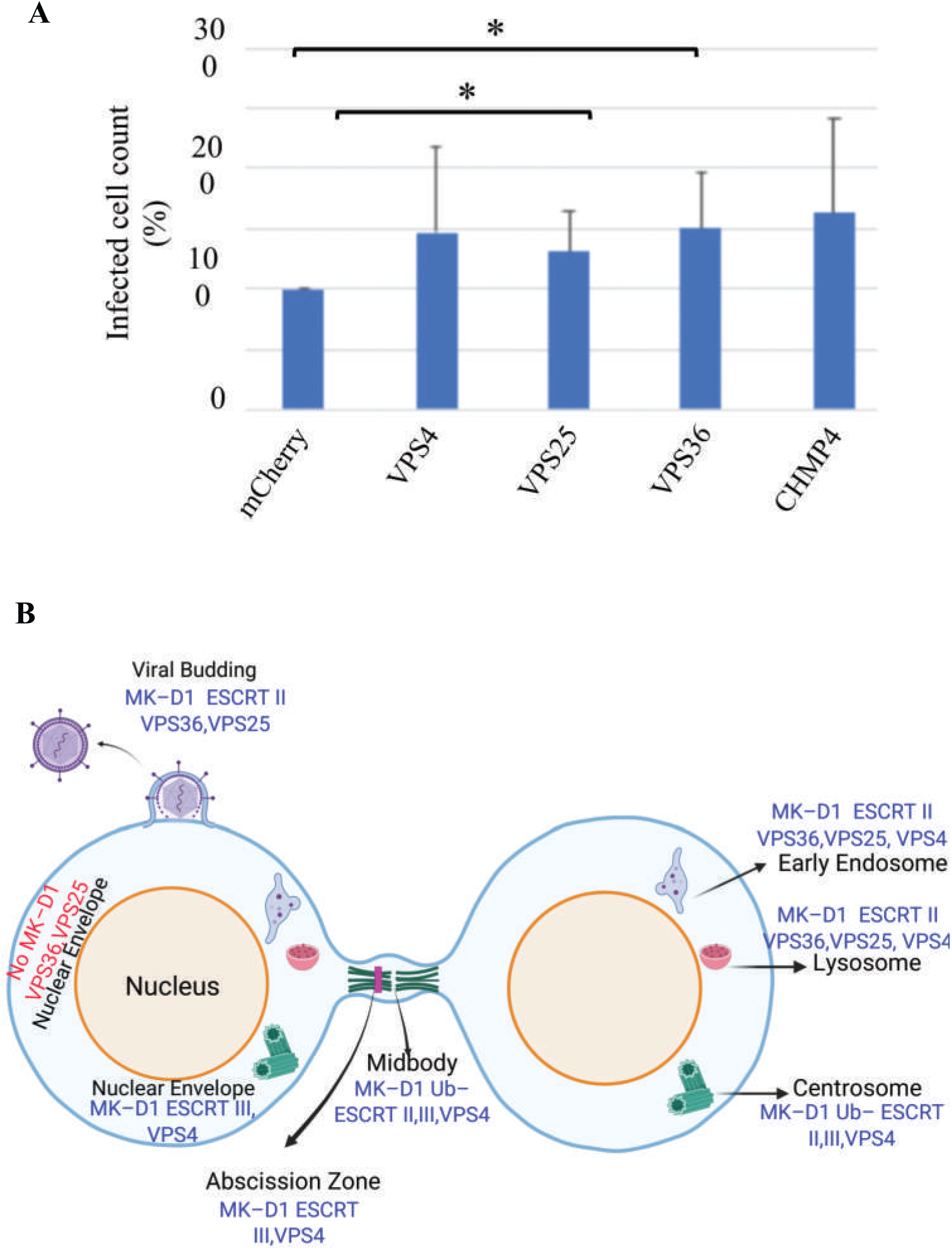
MK-D1 ESCRT modules in viral release and redeployment summary. (A) Retrovirus budding assay. The virus budded from Plat-GP cells expressing mCherry-tagged MK-D1 ESCRT-II/III subunits. The quantity of budded virus was assessed by infecting HEK293 cells. Infection rates increased in viruses budded from cells expressing the respective MK-D1 ESCRT-II/III subunits. However, only increases associated with MK-D1 VPS25 and VPS36 were statistically significant, t-test, n = 6, p = 0.050 and 0.025, respectively; compared to p = 0.134 and 0.076 for VPS4 (n = 6) and CHMP4 (n = 5), respectively. (B) Cartoon showing a summary of MK-D1 ESCRT system redeployment in HeLa cells. Elements are from https://www.biorender.com/.

## Discussion

Our findings provide direct experimental evidence that ESCRT modules from *Candidatus Prometheoarchaeum syntrophicum* (MK-D1), an Asgard archaeon, can integrate into diverse eukaryotic cellular structures and recapitulate features of their eukaryotic counterparts. By deploying archaeal ESCRT-II, ESCRT-III, VPS4, and ubiquitin homologs into human cells, we reveal that these proteins retain the ability to target centrosomes, midbodies, nuclear envelopes, endosomes, and viral budding sites structures that are absent from the archaeal cellular architecture. Despite MK-D1 being bounded by a single membrane, lacking internal organelles or cytoskeletal organizing centers, and not encoding homologs of structural organizers such as CEP55, spastin, or LEM2, its ESCRT proteins displayed remarkable plasticity. The striking localizations highlight a compartment-specific redeployment mechanism, suggesting that the ancestral ESCRT system was modular and capable of being reprogrammed to support the emergence of novel eukaryotic organelles.

Localization of MK-D1 proteins to membrane remodeling sites is consistent with “inside-out” ESCRT activity membrane deformation away from the cytoplasm, as observed in cytokinetic abscission, endosomal budding, and nuclear envelope reformation^41^. These activities align with the biology of Asgard archaea, which are bounded by a single membrane and may utilize ESCRT machinery for processes such as cell division, membrane repair, extracellular vesicle release, and symbiont interactions. The ability of MK-D1 ESCRT-III and VPS4 to form midbody rings, associate with potential abscission sites, and interact with nuclear envelope components underscores the evolutionary continuity of membrane fission mechanisms. Their localization with LEM2 and spastin at reforming nuclear envelopes suggests that nuclear compartment stabilization may have relied on redeploying pre-existing ESCRT modules rather than evolving entirely new machinery, supporting a “topological shift” model in which ESCRT activity expanded from the archaeal cell periphery to internal membranes during eukaryogenesis.

MK-D1 UB localizes to midbodies and centrosomes but not endosomes, likely because the OBF domain in VPS36 replaces the canonical ubiquitin-binding module. This indicates that ancestral recruitment was UB-independent, with UB later adapted for endosomal sorting. In line with the deployment model, MK-D1 ESCRT-II proteins also enhanced viral budding by promoting virus release, highlighting the modularity and flexibility of the ancestral ESCRT system in supporting spatial trafficking cues and opportunistic viral exploitation. The conserved recruitment of MK-D1 proteins to eukaryotic internal compartments suggests that during eukaryogenesis, the ancestral ESCRT system was redeployed to function within newly formed internal membranes while maintaining the same topological direction of action.

In contrast, “outside-in” remodeling, membrane invagination into the cytoplasm has not been observed in Asgard cells^41^. However, in vitro reconstitution experiments indicate that both Asgard and eukaryotic ESCRT-III complexes can mediate inward extrusion of eukaryotic membranes^42–44^, implying that under specific conditions, such as changes in membrane composition or curvature stress, the ancestral ESCRT system may have supported internalization events. This capacity could have contributed to early steps in endomembrane biogenesis by enabling the formation of internal vesicles or invaginations from the cell surface.

The breadth and specificity of Asgard ESCRT proteins’ ability to interface with eukaryotic organelles highlight the uniqueness of this system in providing a direct link between archaeal simplicity and eukaryotic complexity. This modularity allows Asgard ESCRTs to function across multiple compartments, demonstrating a principle of evolutionary exaptation in which ancestral molecular components are repurposed to generate new cellular capabilities. Similarly, minimal Asgard actin regulators, such as profilins and gelsolins, show some capacity to modulate mammalian actin filaments^45,46^. In eukaryotes, these regulators expanded into larger networks, including capping protein, twinfilin, cyclase associated protein and formins, allowing organization of actin in multiple processes such as motility, cytokinesis, and vesicle trafficking^47^. Likewise, the Asgard Sec61/OST/TRAP translocon can engage human ribosomes and, though native to the archaeal cellular membrane, are redirected to the endoplasmic reticulum when expressed in HeLa cells^48^. however, the ESCRT system stands out in its comprehensive integration and multi-site functionality.

Taken together, our study reframes Asgard archaeal ESCRTs as active, modular systems capable of integration into diverse membrane remodeling pathways. Their structural conservation, coupled with divergent regulatory domains, reveals a minimalist yet adaptable toolkit that could be repeatedly redeployed across compartments. Collectively, our findings suggest that the emergence of eukaryotic cellular complexity may be driven by the redeployment and modular expansion of ancestral components. By experimentally demonstrating how Asgard ESCRTs were adapted for multi-compartment integration, this study provides a unique window into the evolutionary trajectory bridging the gap between prokaryotic simplicity and the compartmentalized architecture of eukaryotic cells

## Methods

### Phylogenetic analysis, protein production, purification, crystallization, structure determination, model building, and refinement

A codon-optimized gene encoding MK-D1 VPS25 was synthesized (GeneScript), cloned into the pRSF-Duet expression vector (Merck), and transformed into *Escherichia coli* BL21 (DE3) cells for protein expression. Proteins were purified using nickel-nitrilotriacetic acid (Ni-NTA) affinity chromatography (FUJIFILM Wako), followed by cleavage with HRV 3C protease and further purification by size-exclusion chromatography (Bio-Rad) using standard protocols. Final protein samples were exchanged into the crystallization buffer (10 mM Tris-HCl, pH 8.0, 30 mM NaCl) and concentrated to 10 mg ml^−1^ using centrifugal filters with a 10 kDa molecular weight cut-off (Merck). Crystallization screening of purified MK-D1 VPS25 was performed by the sitting-drop vapor diffusion method. Optimized crystals were grown from 0.1 M Tris, pH 8, 20% PEG 6000, 0.2 M ammonium chloride. Protein crystals were harvested, and cryo-cooled in liquid nitrogen for X-ray diffraction. X-ray data were collected at beamline TPS 05A at the NSRRC (National Synchrotron Radiation Research Center, Taiwan) (1.0 Å) and merged and scaled in HKL2000^49^. Molecular replacement and refinement were carried out using an AlphaFold2-generated model in Phenix^50,51^. Structure predictions of other ESCRT subunits were carried out in AlphaFold3^15^. Structure-based protein sequence alignment and Neighbor-Joining phylogenetic tree construction (100 bootstrap steps) was performed in MAFFT^52^.

### Plasmids

Genes encoding MK-D1 (Asgard archaea) and human ESCRT homologues were synthesized (GenScript), subcloned into pEGFP-C1, pmCherry-C1, or pTagBFP-C1 (Clontech), and used for ectopic expression in HeLa cells. For ESCRT-II components, the following constructs were used: MK-D1 EGFP-Vps36, EGFP-Vps25, and EGFP-Vps22, alongside their human counterparts: mCherry-VPS36, mCherry-VPS25, and mCherry-VPS22. For ESCRT-III and VPS4, constructs included MK-D1 EGFP-CHMP4, EGFP-CHMP1A, EGFP-CHMP1B, and EGFP-VPS4, along with their human homologues: mCherry-CHMP4, mCherry-CHMP2, mCherry-CHMP3, and mCherry-VPS4. An EGFP-tagged MK-D1 ubiquitin construct was also employed. For co-localization analysis, the following subcellular markers were used: TagBFP-CEP55 (midbody and centrosome marker), EGFP-EEA1 (early endosome marker), EGFP-LAMP1A (lysosomal membrane marker), EGFP-LEM2, TagBFP-spastin (nuclear envelope markers), TagBFPMKKS (centrosome marker), and EGFP-α-tubulin.

### Cell culture, transfection and confocal microscopy

HeLa cells (RCB0007, provided by the RIKEN BRC through the National BioResource Project of MEXT, Japan) were maintained in Minimum Essential Medium (MEM; Sigma-Aldrich) supplemented with 10% fetal bovine serum (FBS; Nichirei), 2 mM L-glutamine, and 1× penicillin–streptomycin (FUJIFILM Wako) at 37 °C in a humidified incubator with 5% CO_2_. Cells were routinely tested for mycoplasma contamination using a PCR-based detection kit (Takara Bio).

HeLa cells were seeded in 12-well plates containing 18-mm glass coverslips (Matsunami Glass Ind., Ltd.) for confocal microscopy or 18-mm high-precision glass coverslips (Marienfeld) for super-resolution microscopy, one day before transfection, to reach approximately ∼70% confluency. Transient transfections were performed with either 1 μg of plasmid DNA per well using Xfect transfection reagent (Takara Bio) or 500 ng of plasmid DNA per well with Lipofectamine 2000 (Thermo Fisher Scientific) according to the manufacturer’s protocol. Plasmids encoding EGFP- and mCherry-tagged MK-D1 ESCRT-II, ESCRT-III, and VPS4 constructs, along with their human homologues and subcellular markers for the midbody (CEP55), nuclear envelope (LEM2, spastin), and centrosome (MKKS), were co-transfected. Cells were incubated for 24 h, followed by fixation in 4% paraformaldehyde (Nacalai Tesque) in PBS for 15 min at room temperature. Fixed cells were mounted using FluoroKEEPER antifade mounting medium witrh or without DAPI (Nacalai Tesque) and imaged using an FV1200 confocal laser scanning microscope (Olympus). Co-localization analyses were conducted using Fiji/ImageJ (NIH). All images are representative of at least three independent experiments.

### Live-cell Super-resolution Imaging

Live-cell super-resolution imaging was performed using a spinning disk confocal system (Roper Scientific) coupled with a live super-resolution (SR) module (W1 dual-camera and laser ablation module, Simba, MBI, levels 9–10), mounted on a Nikon Eclipse Ti-E inverted microscope equipped with a Perfect Focus System. The microscope was fitted with a 100× oil immersion objective (Plan Apo VC, NA 1.4) and a high-sensitivity sCMOS camera (Photometrics Prime 95B) for rapid image acquisition. System control and image capture were carried out using MetaMorph software (Molecular Devices). HeLa cells were seeded on 35-mm glass-bottom dishes (Ibidi) and co-transfected with expression constructs encoding fluorescently tagged MK-D1 and human ESCRT homologues II and III, and VPS4 along with subcellular markers using Lipofectamine LTX with PLUS reagent (Thermo Fisher Scientific), following the manufacturer’s protocol. Imaging was conducted 24 h post-transfection under live-cell conditions, with cells maintained in phenol red-free DMEM supplemented with 10% FBS and 25 mM HEPES at 37 °C in a stage-top incubator with CO_2_ control. Z-stack images were acquired using low laser power (488 and 561 nm excitation) and fast acquisition settings to minimize photobleaching and phototoxicity. Live SR mode was enabled to achieve enhanced resolution while maintaining temporal fidelity suitable for dynamic protein localization studies. Z-stacks were collected with step sizes of 200–300 nm and processed using Fiji (ImageJ). Maximum intensity projections of Z-stacks were generated and exported as 16-bit TIFF files for analysis. Image drift and alignment were corrected using the StackReg and MultiStackReg plugins. Colocalization and spatiotemporal recruitment of ESCRT components to membrane subdomains such as midbody, nuclear envelope, and endosomal compartments were analyzed using Fiji software.

### Retrovirus budding assay

Human embryonic kidney HEK293T cells (ATCC CRL-3216) were cultured in Dulbecco’s Modified Eagle Medium (DMEM; Gibco) supplemented with 5% fetal bovine serum (FBS; Nichirei) at 37 °C in a humidified incubator with 5% CO_2_. Plat-GP packaging cells (Cell Biolabs) were maintained in DMEM supplemented with 10% FBS and 1% blasticidin S (FUJIFILM Wako) under the same conditions. To generate retroviruses carrying the GFP gene, Plat-GP cells were seeded into 12-well plates at ∼70–80% confluency one day prior to transfection. Cells were co-transfected with the retroviral expression vector pMX-puro-EGFP, the vesicular stomatitis virus G glycoprotein envelope plasmid (pMD2.G), and one of the following plasmids: pmCherry, pmCherry-VPS4, pmCherry-VPS25, pmCherry-VPS36, or pmCherry-CHMP4, using Lipofectamine 2000 (Invitrogen) according to the manufacturer’s protocol. At 48 h post-transfection, the culture medium containing retroviruses was harvested, centrifuged at 3,000 rpm for 10 min to remove cell debris, and filtered through a 0.22 μm PVDF membrane filter (MILLEX-GV, Millipore). Filtered retroviruses were either used immediately for infection or stored at –80 °C in aliquots for future use. To assess the quantity of bedded viruses, HEK293T cells were seeded in 24-well plates one day before infection and incubated with filtered retroviruses at 37 °C in 5% CO_2_. EGFP expression in infected cells was assessed 48–72 h post-transduction using a CytoFLEX flow cytometer (Beckman Coulter). Relative infection rates were scaled relative to virus budded from mCherry expressing cells. Infection rates represent the average of 5 or 6 repeats. The rabbit anti-GAG antibody (ab100970) was purchased from Abcam. All microscopy images are representative of at least three independent experiments.

## Acknowledgments

We thank the technical support and facilities provided by the National Synchrotron Radiation Research Center, a national user facility funded by the Ministry of Science and Technology, Taiwan. We thank Sriram Muthukumar and Andrew W. Holle from the Mechanobiology Institute, National University of Singapore, for their technical expertise and advice. his research was funded by the Moore-Simons Project on the Origin of the Eukaryotic Cell (grant GBMF9743, R.C.R.), the Japan Society for the Promotion of Science (JSPS, grant JP20H00476 and JP22H04985, R.C.R.; grants 18H02664 and 18K19449, T.H.), and JST CREST (grant JPMJCR19S5, R.C.R.) and Takeda Science, Naito and Ohsumi Frontier Science Foundations (T.H.).

## Author Contributions

Conceptualization and design, K.B.N. and R.C.R.; methodology and investigation, K.B.N. with participation from R.C.R. (Structural biology and bioinformatics), Y.S. (constructs, molecular biology, cell culture, imaging), and Y.B.T. and A.B. (super resolution microscopy); virus experiments, H.L. and T.H.; writing—original draft,K.B.N. and R.C.R.; writing—review & editing, all authors; funding acquisition, R.C.R. and T.H.; supervision, T.H., A.B., Y.B.T., and R.C.R.

## Supplemental Information

**Document S1. Supplemental methods, Figures S1–S7 and Tables S1–S2**.

**Data S1. Movie S1**

## References

1. Imachi, H., Nobu, M.K., Nakahara, N., Morono, Y., Ogawara, M., Takaki, Y., Takano, Y., Uematsu, K., Ikuta, T., Ito, M., et al. (2020). Isolation of an archaeon at the prokaryote-eukaryote interface. Nature 577, 519–525. 10.1038/s41586-019-1916-6.

2. Zaremba-Niedzwiedzka, K., Caceres, E.F., Saw, J.H., Backstrom, D., Juzokaite, L., Vancaester, E., Seitz, K.W., Anantharaman, K., Starnawski, P., Kjeldsen, K.U., et al. (2017). Asgard archaea illuminate the origin of eukaryotic cellular complexity. Nature 541, 353–358. 10.1038/nature21031.

3. Liu, Y., Makarova, K.S., Huang, W.-C., Wolf, Y.I., Nikolskaya, A.N., Zhang, X., Cai, M., Zhang, C.-J., Xu, W., Luo, Z., et al. (2021). Expanded diversity of Asgard archaea and their relationships with eukaryotes. Nature 593, 553–557. 10.1038/s41586-021-03494-3.

4. Rodrigues-Oliveira, T., Wollweber, F., Ponce-Toledo, R.I., Xu, J., Rittmann, S.K.-M.R., Klingl, A., Pilhofer, M., and Schleper, C. (2023). Actin cytoskeleton and complex cell architecture in an Asgard archaeon. Nature 613, 332–339. 10.1038/s41586-022-05550-y.

5. Imachi, H., Nobu, M.K., Ishii, S., Hirakata, Y., Ikuta, T., Isaji, Y., Miyata, M., Miyazaki, M., Morono, Y., Murata, K., et al. (2025). Eukaryotes’ closest relatives are internally simple syntrophic archaea. Preprint at bioRxiv, 10.1101/2025.02.26.640444 https://doi.org/10.1101/2025.02.26.640444.

6. Hatano, T., Palani, S., Papatziamou, D., Salzer, R., Souza, D.P., Tamarit, D., Makwana, M., Potter, A., Haig, A., Xu, W., et al. (2022). Asgard archaea shed light on the evolutionary origins of the eukaryotic ubiquitin-ESCRT machinery. Nat Commun 13, 3398. 10.1038/s41467-022-30656-2.

7. Henne, W.M., Buchkovich, N.J., and Emr, S.D. (2011). The ESCRT Pathway. Developmental Cell 21, 77–91. 10.1016/j.devcel.2011.05.015.

8. Schöneberg, J., Lee, I.-H., Iwasa, J.H., and Hurley, J.H. (2017). Reverse-topology membrane scission by the ESCRT proteins. Nat Rev Mol Cell Biol 18, 5–17. 10.1038/nrm.2016.121.

9. Vietri, M., Radulovic, M., and Stenmark, H. (2020). The many functions of ESCRTs. Nat Rev Mol Cell Biol21, 25–42. 10.1038/s41580-019-0177-4.

10. Henne, W.M., Stenmark, H., and Emr, S.D. (2013). Molecular Mechanisms of the Membrane Sculpting ESCRT Pathway. Cold Spring Harb Perspect Biol 5, a016766. 10.1101/cshperspect.a016766.

11. Christ, L., Wenzel, E.M., Liestøl, K., Raiborg, C., Campsteijn, C., and Stenmark, H. (2016). ALIX and ESCRT-I/II function as parallel ESCRT-III recruiters in cytokinetic abscission. Journal of Cell Biology 212, 499–513. 10.1083/jcb.201507009.

12. Carlton, J.G., and Martin-Serrano, J. (2007). Parallels Between Cytokinesis and Retroviral Budding: A Role for the ESCRT Machinery. Science 316, 1908–1912. 10.1126/science.1143422.

13. Morita, E., Colf, L.A., Karren, M.A., Sandrin, V., Rodesch, C.K., and Sundquist, W.I. (2010). Human ESCRT-III and VPS4 proteins are required for centrosome and spindle maintenance. Proceedings of the National Academy of Sciences 107, 12889–12894. 10.1073/pnas.1005938107.

14. Caspi, Y., and Dekker, C. (2018). Dividing the Archaeal Way: The Ancient Cdv Cell-Division Machinery. Front. Microbiol. 9. 10.3389/fmicb.2018.00174.

15. Abramson, J., Adler, J., Dunger, J., Evans, R., Green, T., Pritzel, A., Ronneberger, O., Willmore, L., Ballard, A.J., Bambrick, J., et al. (2024). Accurate structure prediction of biomolecular interactions with AlphaFold 3. Nature 630, 493–500. 10.1038/s41586-024-07487-w.

16. Hirayama, S., Yamazaki, Y., Kitamura, A., Oda, Y., Morito, D., Okawa, K., Kimura, H., Cyr, D.M., Kubota, H., and Nagata, K. (2008). MKKS Is a Centrosome-shuttling Protein Degraded by Disease-causing Mutations via CHIP-mediated Ubiquitination. Mol Biol Cell 19, 899–911. 10.1091/mbc.E07-07-0631.

17. Martinez-Garay, I., Rustom, A., Gerdes, H.-H., and Kutsche, K. (2006). The novel centrosomal associated protein CEP55 is present in the spindle midzone and the midbody. Genomics 87, 243–253. 10.1016/j.ygeno.2005.11.006.

18. Fabbro, M., Zhou, B.-B., Takahashi, M., Sarcevic, B., Lal, P., Graham, M.E., Gabrielli, B.G., Robinson, P.J., Nigg, E.A., Ono, Y., et al. (2005). Cdk1/Erk2- and Plk1-Dependent Phosphorylation of a Centrosome Protein, Cep55, Is Required for Its Recruitment to Midbody and Cytokinesis. Developmental Cell 9, 477–488. 10.1016/j.devcel.2005.09.003.

19. La Torre, M., Burla, R., and Saggio, I. (2024). Preserving Genome Integrity: Unveiling the Roles of ESCRT Machinery. Cells 13, 1307. 10.3390/cells13151307.

20. Ettinger, A.W., Wilsch-Bräuninger, M., Marzesco, A.-M., Bickle, M., Lohmann, A., Maliga, Z., Karbanová, J., Corbeil, D., Hyman, A.A., and Huttner, W.B. (2011). Proliferating versus differentiating stem and cancer cells exhibit distinct midbody-release behaviour. Nat Commun 2, 503. 10.1038/ncomms1511.

21. Ott, C., Nachmias, D., Adar, S., Jarnik, M., Sherman, S., Birnbaum, R.Y., Lippincott-Schwartz, J., and Elia, N. (2018). VPS4 is a dynamic component of the centrosome that regulates centrosome localization of γ-tubulin, centriolar satellite stability and ciliogenesis. Sci Rep 8, 3353. 10.1038/s41598-018-21491-x.

22. Carlton, J.G., Caballe, A., Agromayor, M., Kloc, M., and Martin-Serrano, J. (2012). ESCRT-III Governs the Aurora B–Mediated Abscission Checkpoint Through CHMP4C. Science 336, 220–225. 10.1126/science.1217180.

23. Morita, E., Sandrin, V., Chung, H., Morham, S.G., Gygi, S.P., Rodesch, C.K., and Sundquist, W.I. (2007). Human ESCRT and ALIX proteins interact with proteins of the midbody and function in cytokinesis. The EMBO Journal 26, 4215–4227. 10.1038/sj.emboj.7601850.

24. Goliand, I., Adar-Levor, S., Segal, I., Nachmias, D., Dadosh, T., Kozlov, M.M., and Elia, N. (2018). Resolving ESCRT-III Spirals at the Intercellular Bridge of Dividing Cells Using 3D STORM. Cell Reports 24, 1756–1764. 10.1016/j.celrep.2018.07.051.

25. Guizetti, J., Schermelleh, L., Mäntler, J., Maar, S., Poser, I., Leonhardt, H., Müller-Reichert, T., and Gerlich, D.W. (2011). Cortical Constriction During Abscission Involves Helices of ESCRT-III–Dependent Filaments. Science 331, 1616–1620. 10.1126/science.1201847.

26. Elia, N., Sougrat, R., Spurlin, T.A., Hurley, J.H., and Lippincott-Schwartz, J. (2011). Dynamics of endosomal sorting complex required for transport (ESCRT) machinery during cytokinesis and its role in abscission. Proceedings of the National Academy of Sciences 108, 4846–4851. 10.1073/pnas.1102714108.

27. Olmos, Y., Hodgson, L., Mantell, J., Verkade, P., and Carlton, J.G. (2015). ESCRT-III controls nuclear envelope reformation. Nature 522, 236–239. 10.1038/nature14503.

28. Vietri, M., Schink, K.O., Campsteijn, C., Wegner, C.S., Schultz, S.W., Christ, L., Thoresen, S.B., Brech, A., Raiborg, C., and Stenmark, H. (2015). Spastin and ESCRT-III coordinate mitotic spindle disassembly and nuclear envelope sealing. Nature 522, 231–235. 10.1038/nature14408.

29. Gatta, A.T., Olmos, Y., Stoten, C.L., Chen, Q., Rosenthal, P.B., and Carlton, J.G. (2021). CDK1 controls CHMP7-dependent nuclear envelope reformation. eLife 10, e59999. 10.7554/eLife.59999.

30. Gu, M., LaJoie, D., Chen, O.S., von Appen, A., Ladinsky, M.S., Redd, M.J., Nikolova, L., Bjorkman, P.J., Sundquist, W.I., Ullman, K.S., et al. (2017). LEM2 recruits CHMP7 for ESCRT-mediated nuclear envelope closure in fission yeast and human cells. Proc Natl Acad Sci U S A 114, E2166–E2175. 10.1073/pnas.1613916114.

31. von Appen, A., LaJoie, D., Johnson, I.E., Trnka, M.J., Pick, S.M., Burlingame, A.L., Ullman, K.S., and Frost, A. (2020). LEM2 phase separation promotes ESCRT-mediated nuclear envelope reformation. Nature 582, 115–118. 10.1038/s41586-020-2232-x.

32. Hadarovich, A., Singh, H.R., Ghosh, S., Scheremetjew, M., Rostam, N., Hyman, A.A., and Toth-Petroczy, (2024). PICNIC accurately predicts condensate-forming proteins regardless of their structural disorder across organisms. Nat Commun 15, 10668. 10.1038/s41467-024-55089-x.

33. Teis, D., Saksena, S., and Emr, S.D. (2008). Ordered Assembly of the ESCRT-III Complex on Endosomes Is Required to Sequester Cargo during MVB Formation. Developmental Cell 15, 578–589. 10.1016/j.devcel.2008.08.013.

34. Henne, W.M., Buchkovich, N.J., Zhao, Y., and Emr, S.D. (2012). The Endosomal Sorting Complex ESCRT-II Mediates the Assembly and Architecture of ESCRT-III Helices. Cell 151, 356–371. 10.1016/j.cell.2012.08.039.

35. Wollert, T., and Hurley, J.H. (2010). Molecular mechanism of multivesicular body biogenesis by ESCRT complexes. Nature 464, 864–869. 10.1038/nature08849.

36. Katzmann, D.J., Babst, M., and Emr, S.D. (2001). Ubiquitin-Dependent Sorting into the Multivesicular Body Pathway Requires the Function of a Conserved Endosomal Protein Sorting Complex, ESCRT-I. Cell 106, 145–155. 10.1016/S0092-8674(01)00434-2.

37. Hurley, J.H. (2015). ESCRTs are everywhere. The EMBO Journal 34, 2398–2407. 10.15252/embj.201592484.

38. Martin-Serrano, J. (2007). The Role of Ubiquitin in Retroviral Egress. Traffic 8, 1297–1303. 10.1111/j.1600-0854.2007.00609.x.

39. Votteler, J., and Sundquist, W.I. (2013). Virus Budding and the ESCRT Pathway. Cell Host Microbe 14, 10.1016/j.chom.2013.08.012. https://doi.org/10.1016/j.chom.2013.08.012.

40. Meusser, B., Purfuerst, B., and Luft, F.C. (2021). HIV-1 Gag release from yeast reveals ESCRT interaction with the Gag N-terminal protein region. J Biol Chem 295, 17950–17972. 10.1074/jbc.RA120.014710.

41. McCullough, J., Frost, A., and Sundquist, W.I. (2018). Structures, Functions, and Dynamics of ESCRT-III/Vps4 Membrane Remodeling and Fission Complexes. Annu. Rev. Cell Dev. Biol. 34, 85–109. 10.1146/annurev-cellbio-100616-060600.

42. Souza, D.P., Espadas, J., Chaaban, S., Moody, E.R.R., Hatano, T., Balasubramanian, M., Williams, T.A., Roux, A., and Baum, B. Asgard archaea reveal the conserved principles of ESCRT-III membrane remodeling. Sci Adv 11, eads5255. 10.1126/sciadv.ads5255.

43. Cada, A.K., Pavlin, M.R., Castillo, J.P., Tong, A.B., Larsen, K.P., Ren, X., Yokom, A.L., Tsai, F.-C., Shiah, J.V., Bassereau, P.M., et al. (2022). Friction-driven membrane scission by the human ESCRT-III proteins CHMP1B and IST1. Proceedings of the National Academy of Sciences 119, e2204536119. 10.1073/pnas.2204536119.

44. Melnikov, N., Junglas, B., Halbi, G., Nachmias, D., Zerbib, E., Gueta, N., Upcher, A., Zalk, R., Sachse, C., Bernheim-Groswasser, A., et al. (2025). The Asgard archaeal ESCRT-III system forms helical filamentsand remodels eukaryotic-like membranes. EMBO J 44, 665–681. 10.1038/s44318-024-00346-4.

45. Akil, C., and Robinson, R.C. (2018). Genomes of Asgard archaea encode profilins that regulate actin. Nature 562, 439–443. 10.1038/s41586-018-0548-6.

46. Akıl, C., Tran, L.T., Orhant-Prioux, M., Baskaran, Y., Manser, E., Blanchoin, L., and Robinson, R.C. (2020). Insights into the evolution of regulated actin dynamics via characterization of primitive gelsolin/cofilin proteins from Asgard archaea. Proceedings of the National Academy of Sciences, 202009167. 10.1073/pnas.2009167117.

47. Gunning, P.W., Ghoshdastider, U., Whitaker, S., Popp, D., and Robinson, R.C. (2015). The evolution of compositionally and functionally distinct actin filaments. J. Cell Sci. 128, 2009–2019. 10.1242/jcs.165563.

48. Carilo, I., Senju, Y., Yokoyama, T., and Robinson, R.C. (2024). Intercompatibility of eukaryotic and Asgard archaea ribosome-translocon machineries. Journal of Biological Chemistry 300. 10.1016/j.jbc.2024.107673.

49. Otwinowski, Z., and Minor, W. (1997). Processing of X-ray diffraction data collected in oscillation mode. Methods Enzymol. 276, 307–326.

50. Jumper, J., Evans, R., Pritzel, A., Green, T., Figurnov, M., Ronneberger, O., Tunyasuvunakool, K., Bates, R., Žídek, A., Potapenko, A., et al. (2021). Highly accurate protein structure prediction with AlphaFold. Nature 596, 583–589. 10.1038/s41586-021-03819-2.

51. Adams, P.D., Afonine, P.V., Bunkoczi, G., Chen, V.B., Echols, N., Headd, J.J., Hung, L.W., Jain, S., Kapral, G.J., Grosse Kunstleve, R.W., et al. (2011). The Phenix software for automated determination of macromolecular structures. Methods 55, 94–106. 10.1016/j.ymeth.2011.07.005.

52. Katoh, K., Rozewicki, J., and Yamada, K.D. (2019). MAFFT online service: multiple sequence alignment, interactive sequence choice and visualization. Briefings in Bioinformatics 20, 1160–1166. 10.1093/bib/bbx108.

