## Supplementary for "Redeployment of Archaeal ESCRT Systems Reveals Conserved Mechanisms Underlying Eukaryotic Complexity"

**Figures**

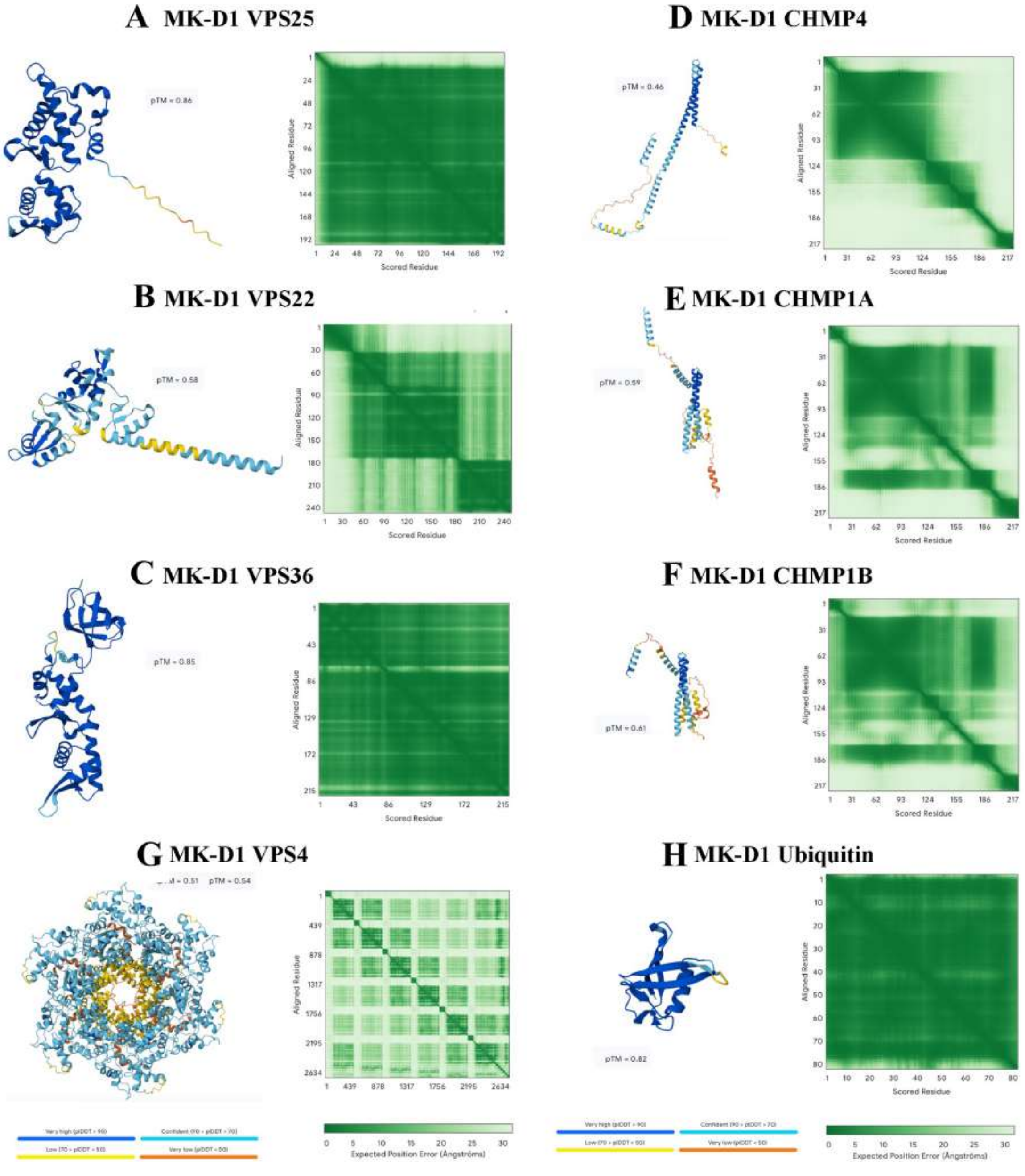


**Figure. S1.** AF3 structure predictions of MK-D1 ESCRT-II/III proteins. In general, MK-D1 ESCRT-II subunits, VPS4, and ubiquitin were well predicted, as indicated by high LDDT (dark blue) and low PAE (dark green) scores. ESCRT-III proteins were predicted with high confidence in their helix–turn–helix motifs.

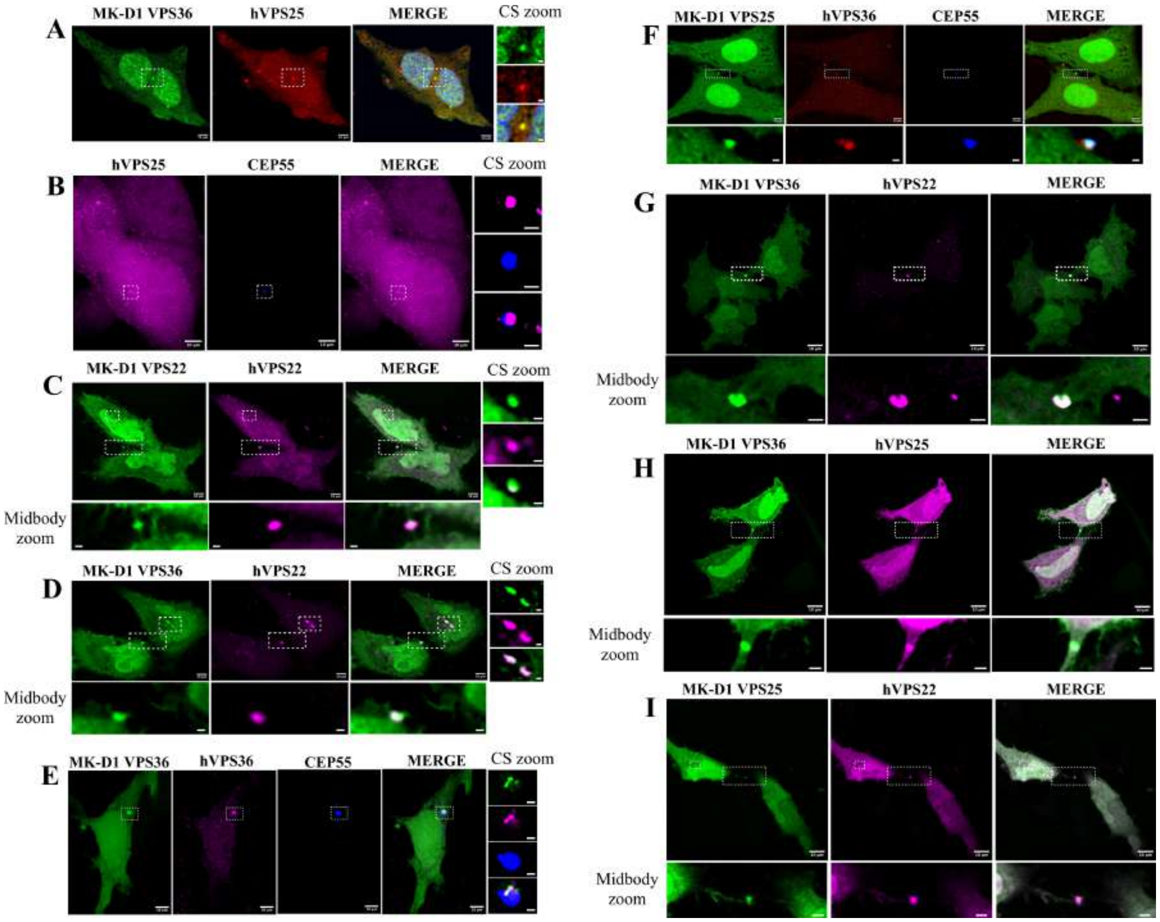


**Figure. S2.** MK-D1 ESCRT-II homologues localize to centrosomes and midbodies. Representative confocal microscopy images of HeLa cells co-transfected with fluorescently tagged MK-D1 and human ESCRT-II constructs.Centrosome (CS): (A) EGFP-MK-D1 VPS36 and mCherry-hVPS25; (B) mCherry-hVPS25 and centrosomal marker TagBFP-CEP55; (C) EGFP-MK-D1 VPS22 and mCherry-hVPS22; (D) EGFP-MK-D1 VPS36 and mCherry-hVPS22; (E) EGFP-MK-D1 VPS36, mCherry-hVPS36 and TagBFP-CEP55. Midbody: (F) EGFP-MK-D1 VPS25 co-localizing with mCherry-hVPS36 and TagBFP-CEP55 at the midbody; (G) EGFP-MK-D1 VPS36 and mCherry-hVPS22; and (H) EGFP-MK-D1 VPS36 and mCherry-hVPS25; (I) EGFP-MK-D1 VPS25 and mCherry-hVPS22.

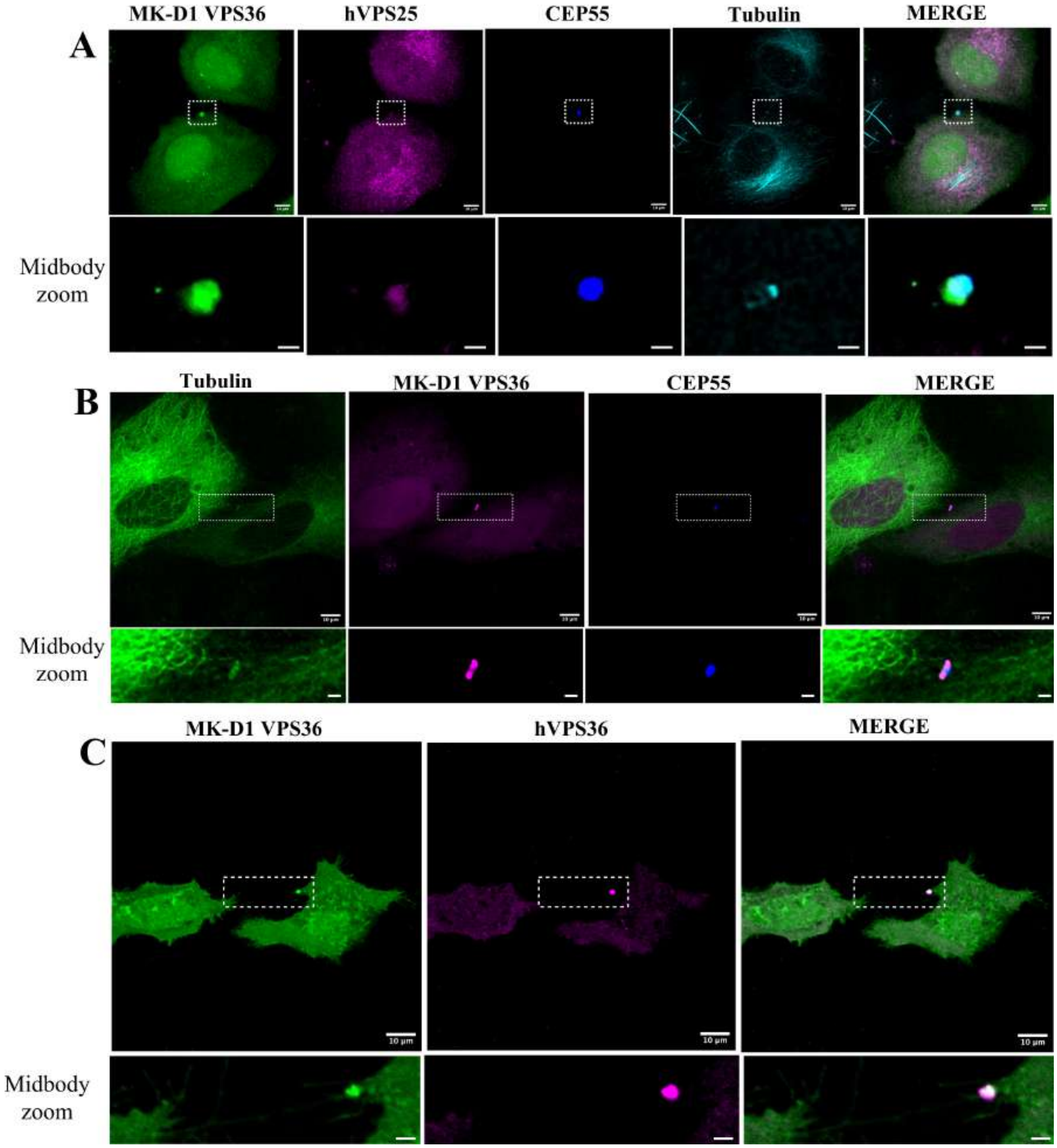


**Figure. S3.** MK-D1 ESCRT-II homologues localize to midbodies. (A) EGFP-MK-D1 VPS36, mCherry-hVPS25, TagBFP-CEP55 and SPY650 tubulin. (B) EGFP-$\alpha$-tubulin, mCherry-MK-D1 VPS36 and TagBFP-CEP55. (C) EGFP-MK-D1 VPS36 and mCherry-hVPS36 indicating midbody inheritance to one daughter cell Scale bars: main images 10 µm, zoom images 2 µm.

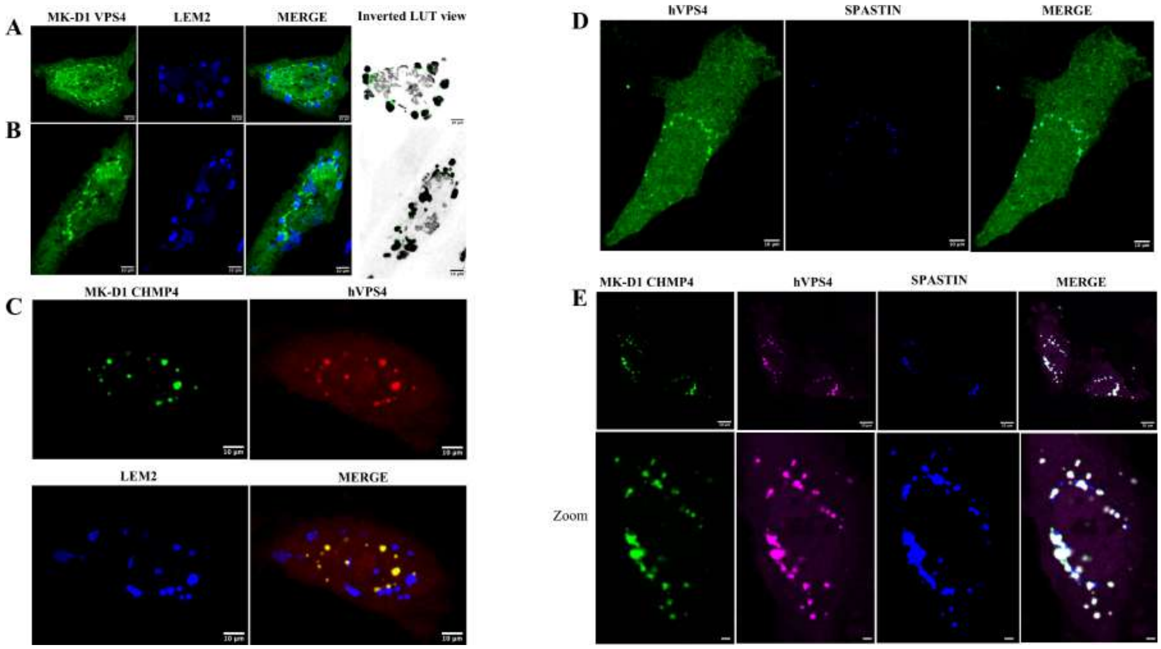


**Figure. S4.** MK-D1 ESCRT-III proteins and VPS4 localize to the reforming nuclear envelope during mitotic exit. Confocal microscopy of HeLa cells co-transfected with fluorescent MK-D1 and human ESCRT-III/VPS4 constructs. (A-B) EGFP-MK-D1 VPS4 and the inner nuclear membrane marker TagBFP-LEM2. (C) EGP-MK-D1 CHMP4, mCherry-hVPS4 and TagBFP-LEM2. (D) EGFP-hVPS4 and TagBFP-spastin. (E) EGFP-MK-D1 CHMP4, mCherry-hVPS4 and TagBFP-spastin. Scale bars: main images 10 μm, zoom images 2 μm.

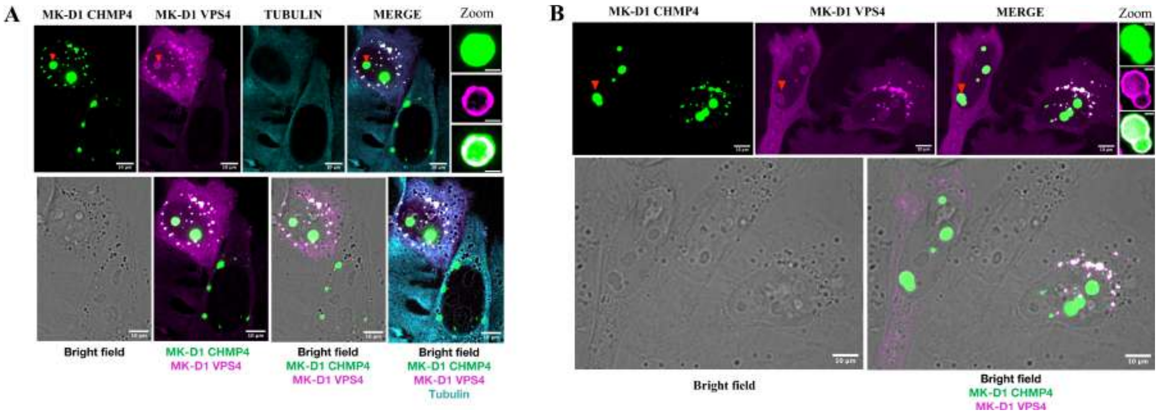


**Figure. S5.** Subcellular organization of MK-D1 CHMP4 and VPS4 foci at nuclear envelope–associated and nuclear-localized structures. (A-B) Super-resolution microscopy. (A) EGFP-MK-D1 CHMP4, mCherry-MK-D1 VPS4 and SPY650 tubulin. (B) EGFP-MK-D1 CHMP4 and mCherry-MK-D1 VPS4. Scale bar: 10 µm. MK-D1 CHMP4 and VPS4 form spherical droplets at the nuclear periphery during mitotic exit, consistent with liquid–liquid phase separation. These cytoplasmic nuclear envelope–associated droplets show uniform distributions of MK-D1 CHMP4 and VPS4, and in some images MK-D1 CHMP4 appear more peripheral to MK-D1 VPS4 (Figure 5E). Larger foci were occasionally observed within the nucleus; however, the distribution of proteins in these foci differed. In these nuclear-localized foci MK-D1 CHMP4 is centrally enriched and VPS4 is peripheral (red arrowheads), indicating compartmentalization. The nuclear foci are visible in the bright field, suggesting they may not represent classical liquid-liquid phase separated condensates.

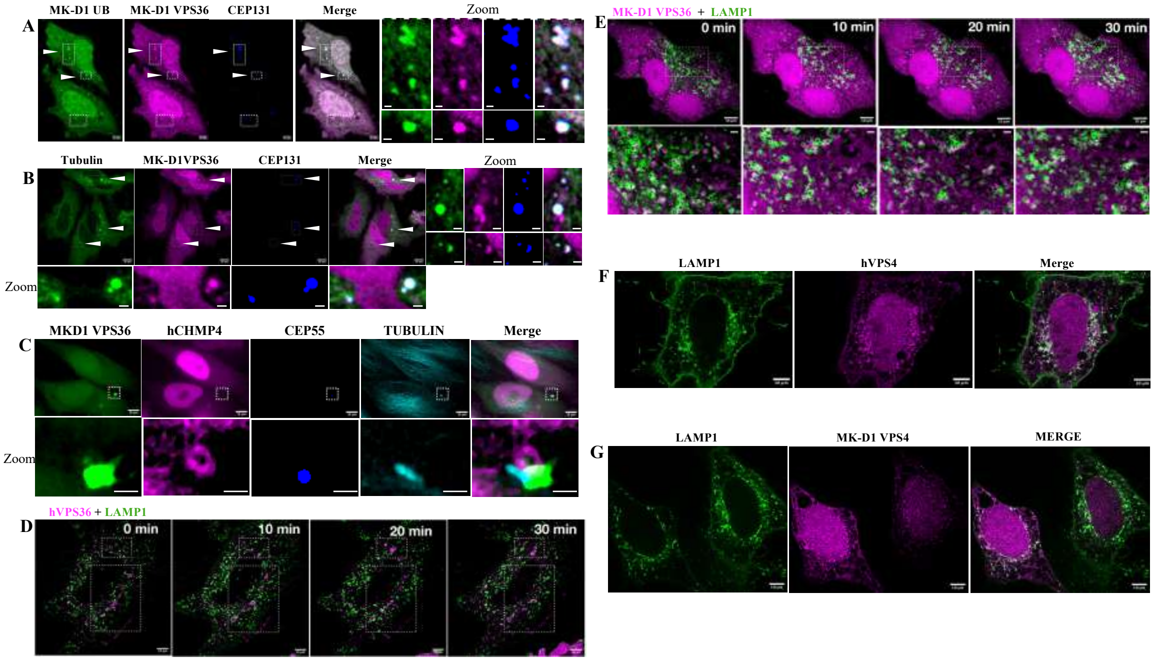


**Figure. S6.** Association of MK-D1 ESCRT, VPS4 and UB with cellular structures. (A) EGFP-MK-D1 UB and mCherry-MK-D1 VPS36 colocalization with the centriolar satellites marker TagBFP-CEP131. (B) EGFP-tubulin and mCherry-MK-D1 VPS36 colocalization with the centriolar satellites marker TagBFP-CEP131. (C) EGFP-MK-D1 VPS36 co-localized with mCherry-hCHMP4, TagBFP-CEP55 and SPY650 tubulin. hCHMP4 forms a ring near proximal to the midbody which overlaps with MK-D1 VPS36. (D-E) Time-lapse imaging of HeLa cells expressing (D) mCherry-hVPS36 or (E) mCherry-MK-D1 VPS36 with EGFP-LAMP1A reveals dynamic reorganization and transient localization of VPS36 with lysosomal compartments (regions highlighted in the boxes). (F-G) mCherry-hVPS4 or mCherry-MK-D1 VPS4 localize to EGFP-LAMP1A positive vesicles. Scale bars: main images 10 μm, zoom images 2 μm.

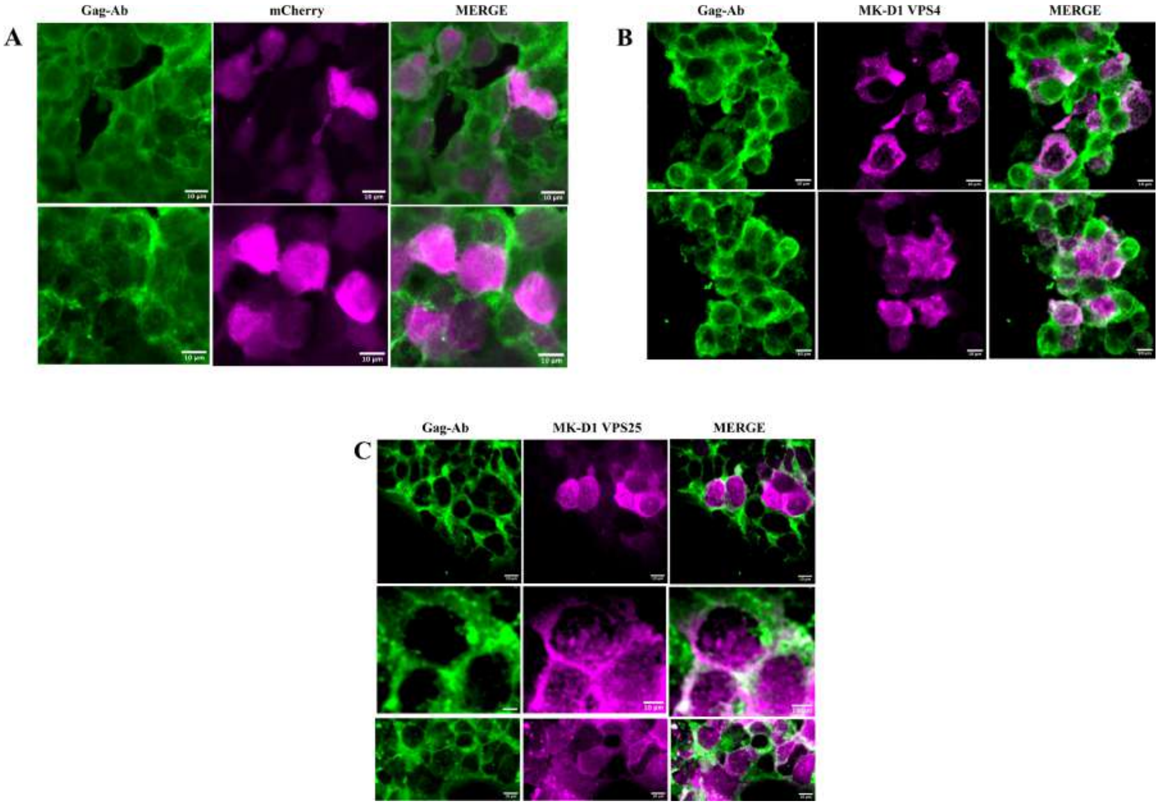


**Figure. S7.** Localization of MK-D1 ESCRT proteins in retrovirus-producing Plat-GP cells. (A) mCherry alone does not co-localize with Gag. (B) mCherry-MK-D1 VPS4 and (C) mCherry-MK-D1 VPS25 show an increased co-localization with Gag, a marker for viral budding sites. Scale bars: 10 µm.

### Tables

| | MK-D1 VPS25<br>QEE17860.1<br>(PDB code 9UYV) |
|---|---|
| **Crystals** | |
| Crystallization Conditions | 0.1 M Tris, pH 8<br>20% PEG 6000<br>0.2 M ammonium chloride |
| Lattice | $P2_1$ |
| *a*, *b*, *c* (Å) | 41.7, 61.1, 88.2 |
| *α, β, γ* (°) | 90.0, 91.1, 90.0 |
| **Data collection** | |
| Beamline | TPS 05A NSRRC |
| Wavelength (Å) | 1.0 |
| Resolution (Å) | 20.0-1.50 (1.53-1.50) |
| $R_{merge}$ | 4.8 (59.8) |
| $R_{meas}$ | 5.6 (72.1) |
| $R_{pim}$ | 2.9 (39.8) |
| *I*/□(*I*) | 25.3 (1.7) |
| $CC_{1/2}$ | (0.723) |
| Completeness (%) | 93.0 (96.9) |
| Redundancy | 3.6 (3.0) |
| **Refinement** | |
| Resolution (Å) | 20.0-1.50 (1.55-1.50) |
| No. reflections | 62930 (4522) |
| $R_{work}$ / $R_{free}$ | 17.4/19.9 (23.6/25.8) |
| No. atoms | |
| Protein | 3120 |
| Water | 688 |
| *B* factors (Å$^2$) | |
| Protein | 18.4 |
| Water | 31.7 |
| r.m.s deviations | |
| Bond lengths (Å) | 0.006 |
| Bond angles (°) | 0.84 |
| Ramachandran Plot | |
| Favoured (%) | 98.0 |
| Outliers (%) | 0.0 |

**Table S1.** X-ray data collection and refinement statistics.

| MK-D1 Protein | PDB | Description | E-Value | Seq. Id. | Rank |
|---|---|---|---|---|---|
| VPS36 C-terminal EAP30 domain | 3CUQ-B | Human VPS36 | 1.91e-6 | 23.0 | 1 |
| | 2ZME-A | Human VPS22 | 2.88e-6 | 22.2 | 3 |
| VPS36 N-terminal oligonucleotide/ oligosaccharide-binding fold | 3WWV-A2 | Pyrococcus horikoshii stomatin operon partner protein | 3.00e-3 | 20.3 | 1 |
| | 8D0K-C | Human CST complex subunit TEN1 | 1.09e-2 | 18.9 | 7 |
| VPS22 C-terminal EAP30 domain | 2ZME-A | Human VPS22 | 4.00e-9 | 20 | 1 |
| | 2ZME-B | Human VPS36 | 1.41e-6 | 15.4 | 4 |
| VPS22 Central DNA-binding domain | 2FU4-A | Escherichia coli Ferric uptake regulation protein | 4.93e-2 | 14.2 | 1 |
| VPS25 structure | 7PB9-A | Odinarchaeota tandem WH domains | 1.44e-10 | 29.4 | 1 |
| | 1XB4-A | Yeast VPS25 | 4.60e-7 | 20.3 | 2 |
| VPS25 AF3 model | 7PB9-A | Odinarchaeota tandem WH domains | 8.58e-11 | 29.6 | 1 |
| | 3CUQ-C | Human VPS25 | 7.04e-7 | 19.7 | 2 |
| CHMP4 | 5T8L-A | Yeast SNF7 | 2.15e-3 | 20.5 | 1 |
| | 6E8G-T | Human CHMP1B | 4.92e-4 | 15.3 | 5 |
| CHMP1A | 6TZ9-A | Human CHMP1B | 3.04e-3 | 19.8 | 1 |
| | 3FRT-B | Human CHMP3 | 2.68e-1 | 16.7 | 6 |
| CHMP1B | 3FRS-A2 | Human IST1 | 4.74e-2 | 16.6 | 1 |
| | 6TZ9-A | Human CHMP1B | 1.54e-2 | 21.6 | 4 |
| VPS4 | 6AP1-C | Yeast VPS4 | 1.00e-26 | 40.3 | 1 |
| | 6P07-E | Drosophila Spastin | 5.21e-25 | 39.1 | 4 |
| Ubiquitin | 7F0N-A | Deaminated human ubiquitin | 3.38e-5 | 23.4 | 2 |
| | 6FJ7-A | Caldiarchaeum Subterraneum Ubiquitin | 8.76e-5 | 19.5 | 15 |

**Table S2.** Closest structural homologs of MK-D1 structures and AlphaFold3 (AF3) models identified in the Protein Data Bank. Structural similarities were determined using FoldSeek[1], with hits ranked by E-value, sequence identity (Seq. Id.), and FoldSeek rank.

**Movie S1 (separate file).** Time-lapse super-resolution imaging of mCherry-MK-D1 VPS36, EGFP-tubulin and TagBFP-CEP55, showing midbody inheritance and internalization into a cell. Movie for Figure. 1.

### SI References


1. van Kempen, M., Kim, S.S., Tumescheit, C., Mirdita, M., Lee, J., Gilchrist, C.L.M., Söding, J., and Steinegger, M. (2023). Fast and accurate protein structure search with Foldseek. Nat Biotechnol, 1–4. https://doi.org/10.1038/s41587-023-01773-0.
